# Site-specific acetylation of polynucleotide kinase 3’-phosphatase (PNKP) regulates its distinct role in DNA repair pathways

**DOI:** 10.1101/2023.06.21.545973

**Authors:** Azharul Islam, Anirban Chakraborty, Altaf H Sarker, Uma K Aryal, Gulshan Sharma, Istvan Boldogh, Tapas Hazra

## Abstract

Mammalian polynucleotide kinase 3’-phosphatase (PNKP) is a dual-function DNA end-processing enzyme with 3’-phosphatase and 5’-kinase activities, which generate 3’-OH and 5’-phosphate termini respectively, as substrates for DNA polymerase and DNA ligase to complete DNA repair. PNKP is thus involved in multiple DNA repair pathways, including base excision (BER), single-strand break (SSBR), and double-strand break repair (DSBR). However, little is known as to how PNKP functions in such diverse repair processes, which involve distinct sets of proteins. In this study, we report that PNKP is acetylated at two lysine (K142 and K226) residues. While K142 (AcK142) is constitutively acetylated by p300, CBP acetylates K226 (AcK226) only after DSB induction. Co-immunoprecipitation analysis using antibodies specific for PNKP peptides containing AcK142 or AcK226 of PNKP showed that AcK142-PNKP associates only with BER/SSBR, and AcK226 PNKP only with DSBR proteins. Although acetylation at these residues did not significantly affect the enzymatic activity of PNKP *in vitro*, cells expressing non-acetylable PNKP (K142R or K226R) accumulated DNA damage, specifically in transcribed genes. Intriguingly, in striatal neuronal cells of a Huntington’s Disease (HD)-based mouse model, K142, but not K226, was acetylated. This observation is consistent with the reported degradation of CBP but not p300 in HD cells. Moreover, genomes of HD cells progressively accumulated DSBs specifically in the transcribed genes. Chromatin-immunoprecipitation analysis using anti-AcK142 or anti-AcK226 antibodies demonstrated an association of Ac-PNKP with the transcribed genes, consistent with PNKP’s role in transcription-coupled repair. Thus, our findings collectively demonstrate that acetylation at two lysine residues located in different domains of PNKP regulates its functionally distinct role in BER/SSBR vs. DSBR.

## Introduction

Mammalian cells continuously incur a plethora of DNA damage induced by various endogenous and exogenous genotoxic agents (1,2). Among myriad types of DNA lesions, DNA strand breaks, [both single strand (SSBs) and double strand (DSBs)] particularly in the transcribed region, pose a major threat to genomic integrity and species survival. Moreover, DNA strand breaks generated in a natural environment, or as DNA base excision repair (BER) intermediates, rarely harbor the canonical 3’-hydroxyl (3’-OH) and 5’-phosphate (5’-P). In most cases, these DNA ends are chemically modified and require further processing. One of the major blocked DNA termini in mammalian cells is the 3’-phosphate (3’-P) and such DNA ends, in addition to impeding DNA repair, can stall elongating RNA polymerases (3). Thus, processing of the “non-ligatable” 3’-P-containing DNA termini is essential for repair progression and efficient transcription in mammalian cells. Another blocked DNA termini is the 5’-OH DNA end, which is generated during Okazaki DNA fragment processing and in some endonuclease-mediated cleavage of genomic DNA (4). Polynucleotide kinase 3’-phosphatase (PNKP) is a bifunctional DNA end-processing enzyme for such blocked DNA termini (3’-P and 5’-OH) at strand breaks in the mammalian genome (5–10). PNKP removes the 3’-P group and catalyzes the phosphorylation of the 5’-OH end to generate the canonical 3’-OH and 5’-P DNA termini, respectively. These termini are necessary for the subsequent activity of DNA polymerase in gap-filling, and DNA ligases in rejoining the two canonical DNA termini and completing the repair process.

Oxidized DNA bases are one of the major endogenous DNA lesions in mammalian cells and are primarily repaired through the Base Excision Repair (BER) pathway. We previously showed that *Nei*-like DNA glycosylase 1 (NEIL1) and NEIL2-initiated repair of oxidized DNA bases predominantly produced 3’-P termini (11,12) which are efficiently processed by PNKP (11,13). We and the Mitra group subsequently established that NEIL1 and NEIL2 preferentially repaired the transcribed genome in mammalian cells via the transcription-coupled BER (TC-BER) pathway where PNKP plays a critical role (14,15). PNKP is the major DNA 3’-phosphatase in mammalian cells (7,16,17) and thus, it is also involved in the repair of SSBs via TC-SSBR. Additionally, several of our recent reports demonstrated the key role of PNKP in nascent RNA-templated error-free DSB repair of 3’P-containing termini in the transcribed genome through the transcription-coupled non-homologous end joining (TC-NHEJ) pathway in mammalian cells (17–19). Our studies further revealed that PNKP associates with RNA polymerase II (RNAPII), various transcription factors, and other repair proteins to form distinct pathway-specific pre-formed repair complexes (15,18,19). However, how PNKP coordinates such multistep and highly complex repair pathways involving distinct sets of proteins remains mostly elusive.

Post-translational modifications of proteins play important roles in diverse cellular processes, including their association with other proteins. It was reported that ionizing radiation (IR) induces phosphorylation of PNKP mediated by Ataxia telangiectasia mutated (ATM) and DNA-dependent protein kinase (DNA-PK), which stimulates its recruitment to DSB sites and modestly enhances the activity of PNKP (20,21). To determine whether PNKP undergoes any additional post-translational modification(s), we affinity purified FLAG-tagged PNKP from stably expressing HEK293 cells and the immunoprecipitated (IP’d) band was analyzed by mass spectrometry (MS). PNKP was indeed found to be acetylated at two sites: constitutively at K142, and at K226 only after the cells were treated with the DSB inducing radiomimetic drug, Bleomycin (Bleo). Notably, unlike other DNA repair proteins, recombinant PNKP, purified from *E. coli*, was found to be acetylated at multiple sites for unknown reasons.

To further validate these acetylation sites, we generated individual cell lines, stably expressing either WT or the mutant (K142R, K226R, and K142R/K226R) PNKP and then purified both WT and the mutant proteins from the corresponding cells (±Bleo) and confirmed by MS analysis that K142 and K226 are the major acetylation sites in PNKP. Finally, we demonstrated that two different acetyl transferases, p300 and CREB-binding protein (CBP), acetylate the two lysines, K142 and K226, respectively. We provided evidence supporting how such distinct site-specific acetylation determines the vital role of PNKP in TC-BER/SSBR vs. TC-NHEJ pathways in mammalian cells.

## Materials and Methods

### Cell culture and treatment conditions

Human embryonic kidney (HEK293) cells and HEK293 stable cell lines, expressing FLAG-tagged WT and mutant (K142R, K226R, K142R/K2226R, K142Q, K226Q and K142Q/K226Q) PNKP, were cultured and maintained in DMEM: F12 (1:1) (Cellgro) containing 10% fetal bovine serum (R & D Systems-Biotechne), 100 units/ml penicillin, streptomycin and amphotericin B (ThermoFisher Scientific) in a 5% CO_2_ incubator at 37 °C with 95% relative humidity. Mouse striatal derived cell line from a knock-in transgenic mouse containing homozygous Huntingtin (HTT) loci with a humanized Exon 1 containing 7 or 111 polyglutamine repeats (Q7 and Q111), respectively, were cultured and maintained in Dulbecco Modified Eagles Medium (high glucose) with 2mM L-glutamine containing 10% fetal bovine serum, 100 units/ml penicillin and streptomycin, and 0.4 mg/ml G418. As Q111 cells lose the ability to proliferate and survive, high-passage cell cultures were avoided. All the cell lines (initial source: ATCC for HEK293 cells and Corriell Institute for the Q7/Q111 cells; Cat# CH00097 and CH00095, respectively) were authenticated by Short Tandem Repeat (STR) analysis in the UTMB Molecular Genomics Core. We routinely tested for mycoplasma contamination in cultured cells using the Mycoalert Mycoplasma Detection Kit (Lonza) according to the manufacturer’s protocol, and the cells were found to be free from mycoplasma contamination. WT and mutant (K/R and K/Q) PNKP expressing stable cell lines were transfected with 10 μM PNKP 3’-UTR specific siRNA (Horizon Discovery) in a 24-well plate using RNAiMAX (ThermoFisher Scientific) according to the manufacturer’s protocol to deplete endogenous PNKP. The sequences of PNKP’s 3’UTR specific siRNA were as follows: sense sequence: CCU CCA CAA UAA ACG CUG UUU, and antisense sequence: ACA GCG UUU AUU GUG GAG G UU. At 48 h post-siRNA transfection, the cells were treated with Glucose oxidase (GO) (400 ng/ml) for 30 min or Bleomycin (Bleo) (30 µg/ml) for 1 h, when cells reached approximately 70% confluency. Likewise, Q7 and Q111 cells were also treated with GO or Bleo under the same experimental conditions.

### Antibodies and Western blotting (WB)

The antibodies (Abs) used in this study and their sources are listed in **Table 1**. WB analysis was performed following the protocol as described earlier (17,19,22,23). All primary Abs were diluted 1:500 in 5% skim milk containing TBST (1X Tris buffered saline with 1% Tween 20) buffer except GAPDH which was diluted to 1:7,000. The dilutions of the secondary HRP-conjugated anti-rabbit and anti-mouse IgG were 1:5,000 and 1:2,000, respectively in the same buffer as mentioned earlier.

**Table 1:** List of antibodies used in this study.

| Primary antibodies | Source | Catalog# | Dilution |
| --- | --- | --- | --- |
| FLAG (M2) | Millipore Sigma | F 1804 | 1:500 |
| Acetyl lysine K142<br>PNKP | Ez Biolabs | Custom-generated | 1:200 |
| Acetyl lysine K142<br>PNKP | Ez Biolabs | Custom-generated | 1:200 |
| PNKP | Bio-Bharati Life<br>Sciences | BB AB0105 | 1:500 |
| CBP | Santa Cruz<br>Biotechnology | Sc-7300 | 1:1000 |
| RNA polymerase II (H5) | Biolegend | 920202 | 1:500 |
| DNA polymerase eta | Cell Signaling<br>Technology | 13848 | 1:500 |
| p300 | Active Motif | 61401 | 1:500 |
| GAPDH | GeneTex | GTX100118 | 1:7000 |
| HDAC2 | GeneTex | GTX109642 | 1:500 |
| XRCC1 | GeneTex | GTX111712 | 1:500 |
| XRCC4 | GeneTex | GTX109632 | 1:500 |
| DNA ligase III | In house (15) |  | 1:500 |
| DNA ligase IV | GeneTex | GTX108820 | 1:500 |
| Ku 70 | GeneTex | GTX101820 | 1:500 |
| Histone H2A.XS139ph<br>(phosphoserine 139) | Cell Signaling<br>Technology | 6731 | 1:500 |
| phospho-53BP1 | Cell Signaling<br>Technology | 2675 | 1:500 |
| 53BP1 | Santa Cruz<br>Biotechnology | Sc-517281 | 1:500 |
| NEIL2 | In house (15) |  | 1:500 |
| DNA polymerase η | Cell signaling<br>Technology | 13848S | 1:500 |
| RNA-DNA hybrid S9.6 | Kerafast | ENH001 | 1:1000 |
| dsDNA Ab | Abcam | ab215896 | 1:2000 |

### Generation of PNKP expressing stable cell lines

For generating the WT and mutant PNKP expressing stable cell lines, HEK293 cells were transfected with PNKP-FLAG (WT and mutants) expressing vector (pcDNA3) in 6-well plates using Lipofectamine 2000 (ThermoFisher Scientific) according to the manufacturer’s protocol. Stably expressing cells were selected with 400 mM geneticin (Millipore-Sigma) starting 48 h post-transfection and continued until the surviving cells formed colonies. The individual colony was further allowed to grow in 9.6 cm^2^ dish with 100 mM geneticin until the cells reached 100% confluency. Finally, WT and mutant PNKP expressing stable cell lines were screened by immunoblotting with anti-FLAG M2 Ab and categorized as low, medium, or high expression cell lines as per the ectopic expression level of PNKP. All other necessary steps were performed as mentioned earlier (22), and as described by the PCDNA3.1 user manual (Thermofisher Scientific).

### Site-directed mutagenesis

The coding DNA sequence (CDS) of the human PNKP (gene accession#NM_007254.4) was amplified using the Q5 hot start high fidelity DNA polymerase (New England Biolabs), with HEK293 genomic DNA as a template, and the amplified PCR fragment was cloned in pCDNA3 (containing an N-terminal FLAG tag) using *Hin*dIII-*Bam*HI sites as described earlier (17). K142 and K226 were mutated to Arginine (K142R & K226R) or glutamine (K142Q & K226Q), individually, using the Q5 site-directed mutagenesis kit (New England Biolabs), according to the manufacturer’s protocol. PCR primers for site-directed mutagenesis were designed using NEBaseChanger. The primers to introduce these mutations are listed in **Table 2**. A double mutant (K142R/K226R) was generated using the single mutant K142R as a template and then the K226R mutation was introduced in it. Similarly, a K142Q/K226Q double mutant was generated using K142Q as a template.

**Table 2:**
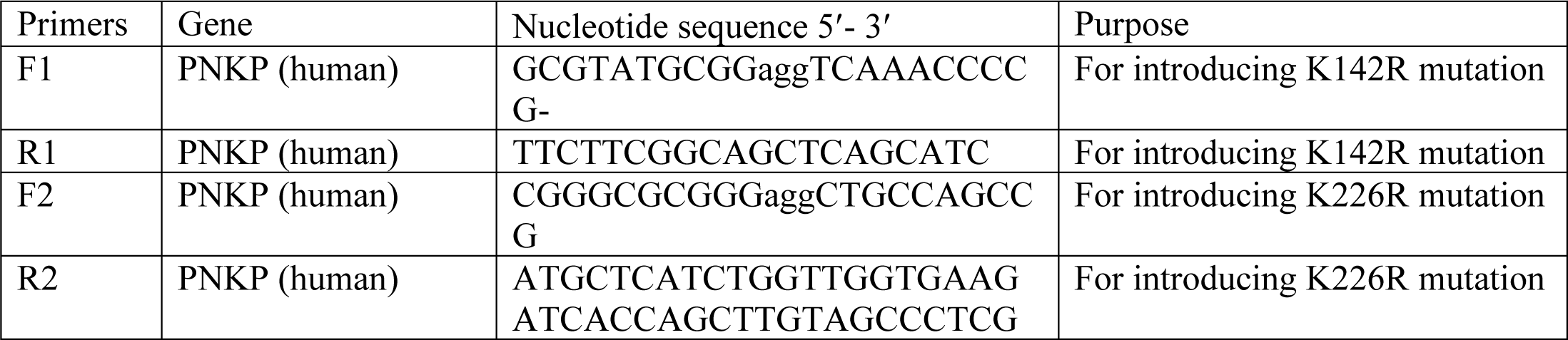
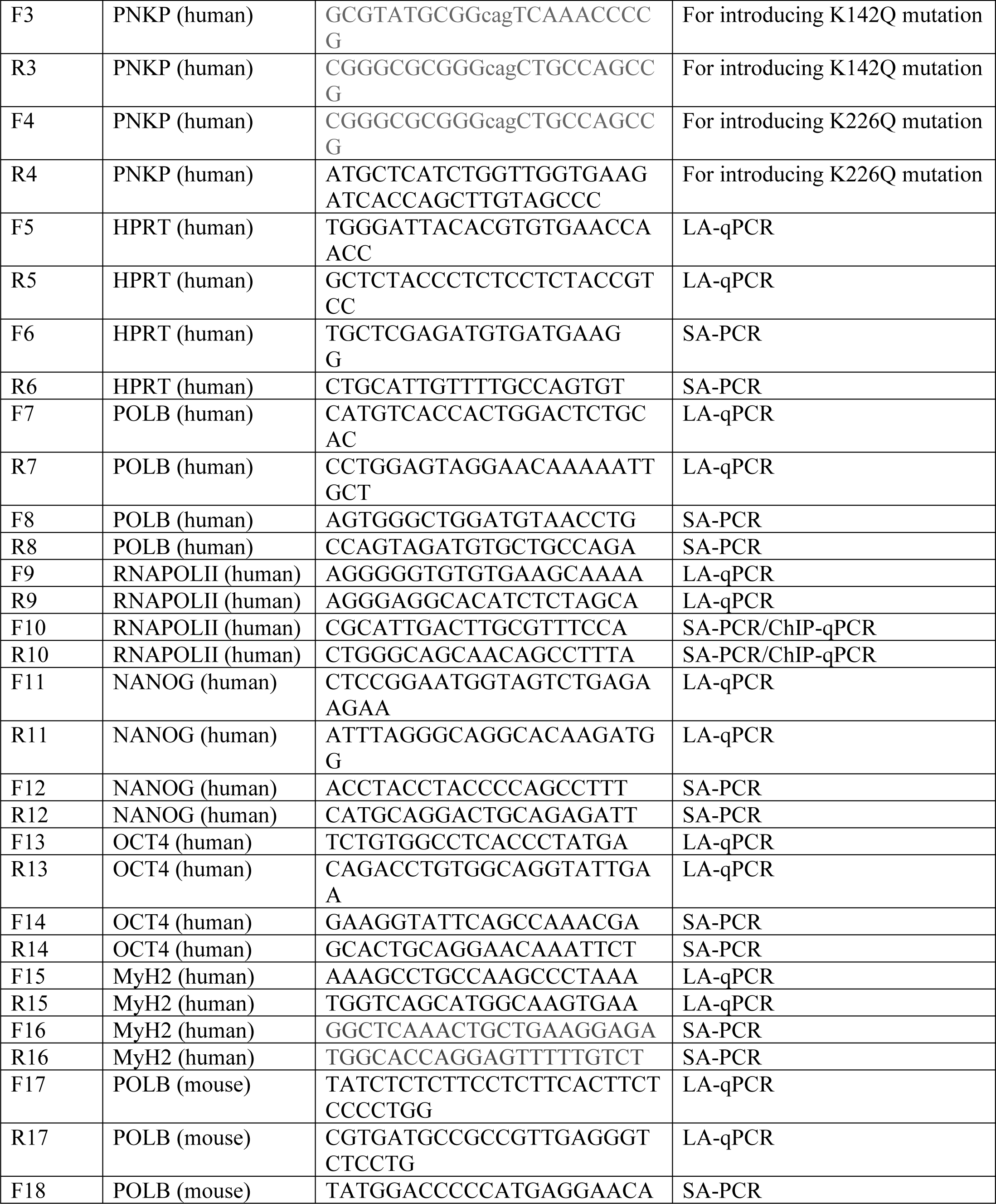
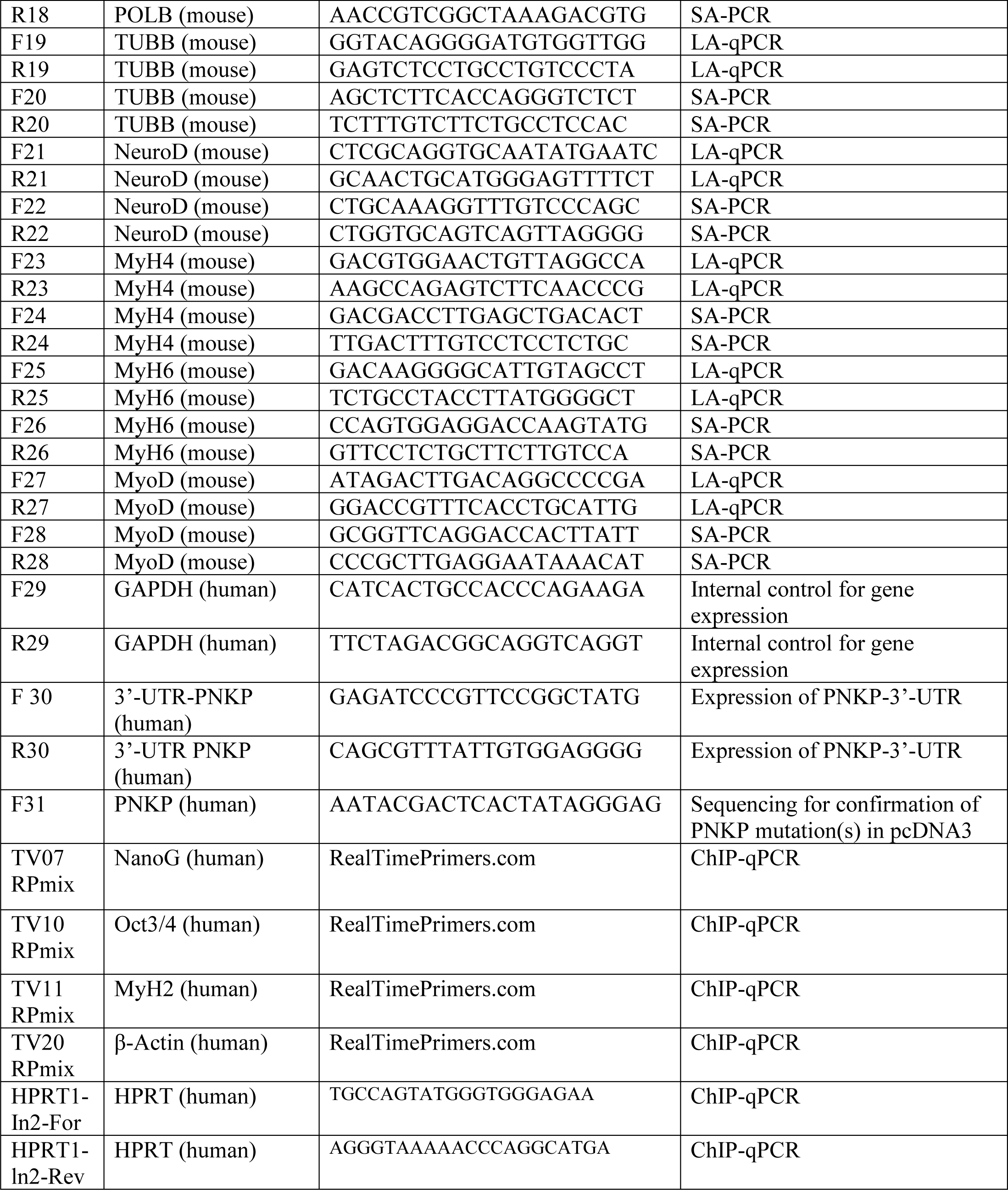
Primers used in the study.

### Cell fractionations and immunoprecipitation (IP)

Cytoplasmic, nuclear, and chromatin fractions were prepared from GO or Bleo-treated WT or mutant cells following the previously described protocol (17,24), with modifications as follows: Dithiothreitol (DTT) was excluded from the cytosolic, nuclear and chromatin fractionation buffers due to interference of DTT with the binding of FLAG-PNKP to Anti-FLAG M2 Affinity gel, which contains a mouse monoclonal Ab that is covalently attached to agarose (Millipore Sigma). Additionally, 0.30 units/μL of benzonase (MilliporeSigma) was added to the chromatin fractionation buffer followed by incubation at 37 °C for 45 min. For IP, 30 μl of Anti-FLAG M2 Affinity gel was mixed with an appropriate volume of chromatin fraction containing 1 mg of total protein, and incubated in a rotatory shaker overnight at 4 °C. The immune complex was washed 3 times with wash buffer (20 mM HEPES pH 7.9, 0.5 mM EDTA, 10% glycerol, 0.25% Triton X-100, 250 mM KCl, 1× cOmplete mini EDTA-free protease inhibitor cocktail (Roche)), and FLAG-PNKP was eluted from the affinity gel using elution buffer (20 mM Tris HCl pH 7.5, 150 mM NaCl, 10% glycerol, protease inhibitor cocktail) containing 150 mM FLAG peptide (Millipore Sigma). 20 μL of the eluted product was separated on SDS-PAGE and stained with Coomassie Brilliant Blue (CBB) followed by destaining, and the concentration of PNKP protein was determined using known standards of BSA. The purified PNKP was used for downstream processes. The concentration of eluted WT and mutant FLAG-PNKP protein was further confirmed by densitometric analyses of immunoblot compared with the affinity-purified recombinant His-PNKP of known concentration, using 3 μL eluted PNKP for SDS-PAGE separation (17,23). Using 10 μg of custom-made rabbit polyclonal anti-AcK142 Ab (Ez Biolabs), AcK142 PNKP was IP’d from 1 mg of GO-treated chromatin fraction from WT-PNKP-FLAG and K142R-PNKP-FLAG expressing cells. Similarly, IP was performed on the chromatin fraction of mock vs. Bleo-treated WT-PNKP-FLAG and K226R-PNKP-FLAG cells using 10 μg of custom-made rabbit polyclonal anti-AcK226 Abs (Ez Biolabs). All other subsequent steps and buffers used for IP were the same as described earlier (17,18). Immunocomplexes were separated in SDS-PAGE and probed with anti-FLAG Ab to detect AcK142 and AcK226 PNKP in the respective samples. Further, Co-IP experiments were performed using anti-AcK142 or anti-AcK226 Abs from chromatin fractions of GO or Bleo-treated WT-PNKP-FLAG expressing cells, respectively, to characterize respective immunocomplexes. The immunocomplexes were tested for the presence of different SSB and DSB repair proteins using appropriate Abs.

### Mass spectrometry analysis by LC-MS/MS

Mass spectrometry analysis by LC-MS/MS was performed at the Taplin Biological Spectrometry facility at Harvard Medical School. IP of PNKP was performed as described in the *Cell Fractionations and IP* section. 30 µL IP’d WT and mutant (K226R) PNKP were separated by SDS-PAGE and gel bands of PNKP were excised after Coomassie Brilliant Blue (CBB) staining. Following destaining, excised gel bands were sliced into approximately 1 mm^3^ pieces. Gel pieces were subjected to a modified in-gel trypsin digestion procedure (25) and the sample was run over a nanoscale processed reverse-phase HPLC capillary column (26). Eluted peptides were subjected to electrospray ionization and then entered an LTQ Orbitrap Velos Pro ion-trap mass spectrometer (Thermo Fisher Scientific). Peptide sequences (and hence protein identity) were determined by matching protein or translated nucleotide databases with the acquired fragmentation pattern by the software program, Sequest (Thermo Finnigan) (27). The differential modification of 42.0106 mass units to lysine was included in the database searches to determine acetylated peptides. All databases include a reversed version of all sequences, and the data were filtered to a 1-2% peptide false discovery rate.

### Microscopic imaging

WT and mutant (K-R) PNKP expressing stable cell lines were transfected with the PNKP 3’-UTR specific siRNA to deplete endogenous PNKP (22) as described above. Cells (1×10^5^) were then trypsinized and plated on collagen pre-treated cover glasses (Roche Applied Sciences) followed by treatment with GO or Bleo. Likewise, Q7 and Q111 cells were also treated with GO or Bleo. Cells were then fixed with an acetone-methanol (1:1) solvent for 10 min at room temperature, and dried. Next, cells were rinsed and permeabilized using 0.1% (w/v) Tween-20 diluted in phosphate-buffered saline (PBST) for 5 min, incubated with 1% BSA for 1 h at room temperature, and then blocked with anti-mouse IgG (1:100 dilution) for 1 h at 37 °C. WT (Q7) and Huntington’s Disease (HD)-derived mouse striatal neuronal cells (Q111) were treated with 1% Triton X-100 for 15 min at room temperature, instead of Tween-20, for efficient permeabilization. Cells were incubated with custom-generated anti-AcK142 and anti-AcK226 Abs (EZ BioLabs) at a dilution of 1:200 in PBST for 1 h at 37 ⁰C. After 3 washes in PBST, the cells were incubated with a secondary Ab conjugated to Alexa Fluor 594 (goat anti-rabbit) for 1 h at 37 ⁰C. After washing in PBST (3 times), cells were dried and mounted with Vecta shield/DAPI, 4′6-diamidino-2-phenylindole hydrochloride (Vector Laboratories). More than 20 randomly selected fields of view per sample were photographed using a WHN10×/22 eyepiece and a 60× objective (field of view is 1.1 mm, and camera correction is 1.0) on an Echo Revolution Microscope system.

### *In vitro* 3’-phosphatase and 5’-kinase assays of PNKP

The phosphatase activity of IP’d PNKP from mock vs. Bleo-treated WT, K142R, K226R, and K142R/K226R cells was determined by the *in vitro* 3’-phosphatase assay as described earlier (13,16,18,22) with minor modifications. Briefly, 1 ng of IP’d WT or mutant PNKP was incubated with γP^32^ ATP radiolabeled 3’-phosphate-containing SSB-mimic substrate (5 pmole) for 13 min at 37 °C in phosphatase assay buffer (25 mM Tris-HCl pH 7.5, 100 mM NaCl, 5 mM MgCl_2_, 1 mM DTT, 10% glycerol, and 0.1 μg/μL acetylated BSA). Additionally, the 5’-kinase activity assay with the IP’d WT vs. mutant PNKP was essentially performed as previously described (28–30) with some modifications. Briefly, γP^32^ labeled ATP was incubated in kinase assay buffer (80 mM succinic acid pH 5.5, 10 mM MgCl_2_, 1 mM DTT, 2.5% glycerol) along with 1.0 μg/μl acetylated BSA, and 0.6 pmole labeled substrate for 30 min at 30 °C. 100 fmol of PNKP and 2.5 pmole of cold substrate were used in this assay. For both the 3’-phosphatase and the 5’-kinase assays, the radioactive bands were visualized using a PhosphorImager (GE Healthcare) and quantitated using ImageQuant software. The data were represented as % product released from the radiolabeled substrate.

### Reverse transcription PCR

Control or PNKP 3’UTR siRNA-transfected and mock or GO/Bleo treated cells were used to assess the depletion of endogenous PNKP by RT-PCR. Total RNA was extracted using TRIzol (ThermoFisher Scientific) according to the manufacturer’s protocol. 1 μg of total RNA was used to make cDNA libraries using prime script RT reagent kit gDNA eraser (Takara Bio Inc.) according to the kit’s protocol, and RT-PCR was carried out using 1 μl of cDNA to amplify the 3’UTR region of the endogenous PNKP gene, as well as the housekeeping gene (GAPDH) using quick-load Taq 2× master mix (New England Biolabs), according to manufacturer’s protocol. The primers used for amplifying the 3’UTR region of PNKP and GAPDH are listed in **Table 2**. The amplified PCR products were run in 1.5% agarose gel and stained with ethidium bromide, and the image was processed by Gel Doc EZ imager (BioRad). The quantitation of gel bands was performed using ImageJ software and the endogenous PNKP expression was normalized using GAPDH. The relative expression levels were presented with the expression of endogenous PNKP in control siRNA transfected cells considered as 100.

### Long amplicon quantitative PCR (LA-qPCR)

Extraction of genomic DNA from WT, K-R or K-Q mutant cells (post PNKP 3’-UTR siRNA transfection and GO/Bleo treatment) was performed using the QiaAmp DNA Micro kit (Qiagen) according to the manufacturer’s protocol, with the concentration determined by the NanoVue (GE Healthcare). Gene-specific LA-qPCR assays for measuring DNA strand-breaks (SBs) were performed as described earlier (7,17,18,22) using Long Amp Taq DNA Polymerase (New England BioLabs). 10 ng of genomic DNA was used as a template to amplify transcribed genes (10.4 kb region of the hypoxanthine-guanine phosphoribosyltransferase [HPRT], 12.2 kb of the DNA polymerase beta [Pol Beta], 11.3 kb of the RNA polymerase II [RNA pol II] and non-transcribed (8.6 kb of the Nanog homeobox [NANOG], 10.1 kb of the POU class 5 homeobox 1 [OCT 4] and 6.0 kb of the myosin heavy polypeptide 2 [MYH2]), using the primers described previously (**Table 2**)(7,17,18). Similarly, LA-qPCR was performed from genomic DNA isolated from Q7 and Q111 cells. Since these are neuronal cells, a different set of transcribed (neuronal differentiation factor 1 (NeuroD), tubulin β3 class III (TUBB) and POLB) vs. non-transcribed (myogenic differentiation factor [MyoD], muscle-specific myosin heavy chain 4 and 6 [MyH4, MyH6]) genes were used for the LA-qPCR assay. Since a short region would have less probability of DNA SBs, a short DNA fragment (short amplicon PCR; SA-PCR) from the same gene was also amplified for normalization of the amplified long fragment. To overcome the variation in PCR amplification while preparing the PCR reaction mixture, the LA-qPCR and SA-PCR reactions were set for all genes from the same stock of diluted genomic DNA samples. The amplified products were separated in 0.8% agarose gel and then stained with ethidium bromide followed by imaging by Gel Doc EZ imager (BioRad). The band intensity was determined by ImageJ software and the extent of DNA damage was calculated based on the relative quantification of the ratio of the long fragment and short fragment.

### Lactate dehydrogenase (LDH) cytotoxicity assay

HEK293 cells expressing WT, K142R, K226R and K142R/K226R PNKP were counted after trypsinization, and 1×10^4^ cells were plated in a 24-well plate. Cells were transfected with PNKP 3’ UTR specific siRNA for the depletion of endogenous PNKP and then were individually treated with Bleo for 1 h, at 48 h post-transfection of siRNA. Subsequently, the LDH cytotoxicity assay was performed according to the manufacturer’s protocol (BioLegend, cat#426401). The colorimetric reading was recorded immediately at OD 450 nm on a microplate reader (Synergy H1 Hybrid Multi-Mode Reader; BioTek) at 37 °C under light-protected conditions. The amount of released LDH in media due to Bleo-induced DNA double-strand breaks was determined using the formula ΔA450 nm = (A2 – A1), where: A1 is the sample reading at 0 h and A2 is the sample reading at 2 h. DMEM without phenol red was used as a background control in this assay. LDH activity was expressed as nmoles of NADH generated by LDH reaction during the reaction time (ΔT = T2 – T1). LDH activity was determined using the formula (BI (ΔTx V)) × D in nmol/min/mL units, where B = amount of NADH in the sample, calculated from a standard curve (nmol); ΔT = Reaction time (minutes); V = Original sample volume added into the reaction well (mL); and D = Sample dilution factor.

### Chromatin immunoprecipitation (ChIP) assay

ChIP assays were performed essentially as described earlier (17,31) with minor modifications. Briefly, mock, GO or Bleo-treated WT-PNKP FLAG cells were incubated with 1% formaldehyde for 15 min at room temperature for cross-linking. Subsequently, the genomic DNA was fragmented to a size range of 300-500 bp by sonication (3-5 pulses of 30 seconds at 30 amplitude) using a sonicator (Qsonica-Q700). Total protein concentration was determined by a Bradford protein assay, and 100-500 μg of protein lysate was used for each IP. 10 μg of ChIP-grade, custom-made rabbit polyclonal anti-AcK142 or anti-AcK226 (Ez Biolabs) Abs were incubated overnight at 4 °C with the lysate from mock vs. GO-treated or mock vs. Bleo-treated cells, respectively. Ab-protein-DNA complexes were captured by protein A-G agarose beads (Santa Cruz Biotechnology) for 4 h at 4 °C. Buffers used and all subsequent steps including the elution of the immune complex, de-crosslinking, digestion of proteins and purification of the DNA were performed following the procedures as described earlier (17,31). The purified DNA was subjected to qPCR using CDS-specific primers (transcribed and non-transcribed gene-specific) as listed in **Table 2**.

### Statistical analysis

A two-sided unpaired Student’s *t*-test (http://www.ruf.rice.edu/~bioslabs/tools/stats/ttest.html) was used for the analysis of statistical significance between two sets and among three sets of data. The level of significance was evaluated at levels *P*>0.05 (ns), *P*<0.05 (*), *P*<0.01 (**) and *P*<0.005 or 0.0001 (***), as the case may be.

## Results

### Human PNKP is acetylated at amino acid residues K142 and K226

The role of phosphorylation in modulating the DNA repair function of PNKP was previously investigated (21,32). To identify additional post-translational modification in PNKP, FLAG-tagged PNKP was affinity-purified from stably expressing HEK293 cells, either mock-treated or treated with the radiomimetic drug, Bleomycin, to induce DSBs. The affinity-purified PNKP was then analyzed by mass spectrometry. PNKP was found to undergo post-translational modification via acetylation at two different lysine residues (K142 and K226), as indicated in bold in the peptide fragments, R/**K142**SNPGWENLEK/L and R/G**K226**LPAEEFK/A (**Supplementary** Figs. 1A; **1B and 1C**). K142 acetylation (AcK142) was found to be constitutive and identified in both control and Bleo-treated cells, whereas K226 acetylation (AcK226) was induced only upon Bleo treatment. These two distinct acetylation sites were found to be located in different domains of PNKP (6,30,33,34): AcK142 in the linker region (111-144 aa), and AcK226 in the phosphatase domain (145-336 aa) (**Supplementary** Fig. 1D). To validate the acetylation sites in PNKP and the differential mode of their acetylation, we further performed the MS analysis of affinity-purified FLAG-tagged PNKP from WT and K226R (acetylation-deficient mutant) PNKP expressing HEK293 cells following mock and Bleo-treatment. We observed acetylation of PNKP at K142 in both control and Bleo-treated WT and K226R cells. However, acetylation at K226 was observed in the WT cells only following Bleo-treatment and no such acetylation was observed in the K226R mutant cells (**Supplementary** Fig. 1E). These data further confirmed two distinct acetylation sites in PNKP, of which K142 is acetylated constitutively, whereas K226 is acetylated upon DSB induction.

### Site-specific acetylated lysine antibodies validated the acetylation of PNKP at K142 and K226

To confirm MS-identified PNKP acetylation sites, we custom-generated PNKP acetylation site-specific Abs against individual peptide fragments containing acetylated K142 or K226. We then validated such antibodies for target specificity by IP of acetylated PNKP from the chromatin fraction of FLAG-tagged WT and acetylation-deficient (K142R or K226R) PNKP expressing HEK293 cells ± Glucose oxidase (GO, SSB inducing agent) or Bleomycin treatment. The IP of PNKP with anti-AcK142 Ab followed by the detection of the IP’d band using anti-FLAG Ab through Western analysis, showed that acetylation at K142 occurred only in WT PNKP expressing but not in GO-treated K142R cells (**Fig. 1A**). Indirect immunofluorescence analysis of the individual cell population using anti-AcK142 Ab demonstrated the nuclear localization of AcK142 PNKP in WT (**Fig. 1B, top panel)** and K226R (**Fig. 1B, bottom panel)** cells, but not in the K142R (**Fig. 1B, middle panel**) expressing cells. Therefore, both Western analysis and microscopic imaging data identified the K142-specific constitutive acetylation in PNKP. Similarly, IP of PNKP with anti-AcK226 Ab showed that acetylation at K226 occurred only in the Bleo-treated WT cells (**Fig. 1C, lane 3, upper panel)**. Neither the mock (**Fig. 1C, WT, lane 1 and K226R, lane 2)** nor the Bleo-treated K226R (**Fig. 1C, lane 4)** cells showed the presence of AcK226. Immunofluorescence analysis also revealed that K226 of PNKP is acetylated only in Bleo-treated (**Fig. 1D, 2nd panel from the top)**, but not in mock-treated WT (**Fig. 1D, top panel)**, nor in mock (**Fig. 1D, 3rd panel from the top)** or Bleo-treated K226R (**Fig. 1D, bottom panel)** cells. These Abs did not detect nor cross-react with unmodified PNKP, thereby confirming the specificity of affinity-purified antibody. We further confirmed the presence of DSB markers, phospho-p53-binding protein 1 (p-53BP1) **(Supplementary** Fig. 2A**, lanes 1-2, top panel)** and γH2AX **(Supplementary** Fig. 2A**, lanes 1-2, 3rd panel from the top)** in Bleo-treated WT and K226R cells but not in GO-treated WT and K142R **(Supplementary** Fig. 2A**, lanes 3-4)** cells. Detection of acetylation by indirect immunofluorescence was performed following the depletion of endogenous PNKP by 3’-UTR specific siRNA, thereby excluding the possibility of any interference of acetylation from endogenous PNKP **(Supplementary** Fig. 2B).

**Figure 1.**
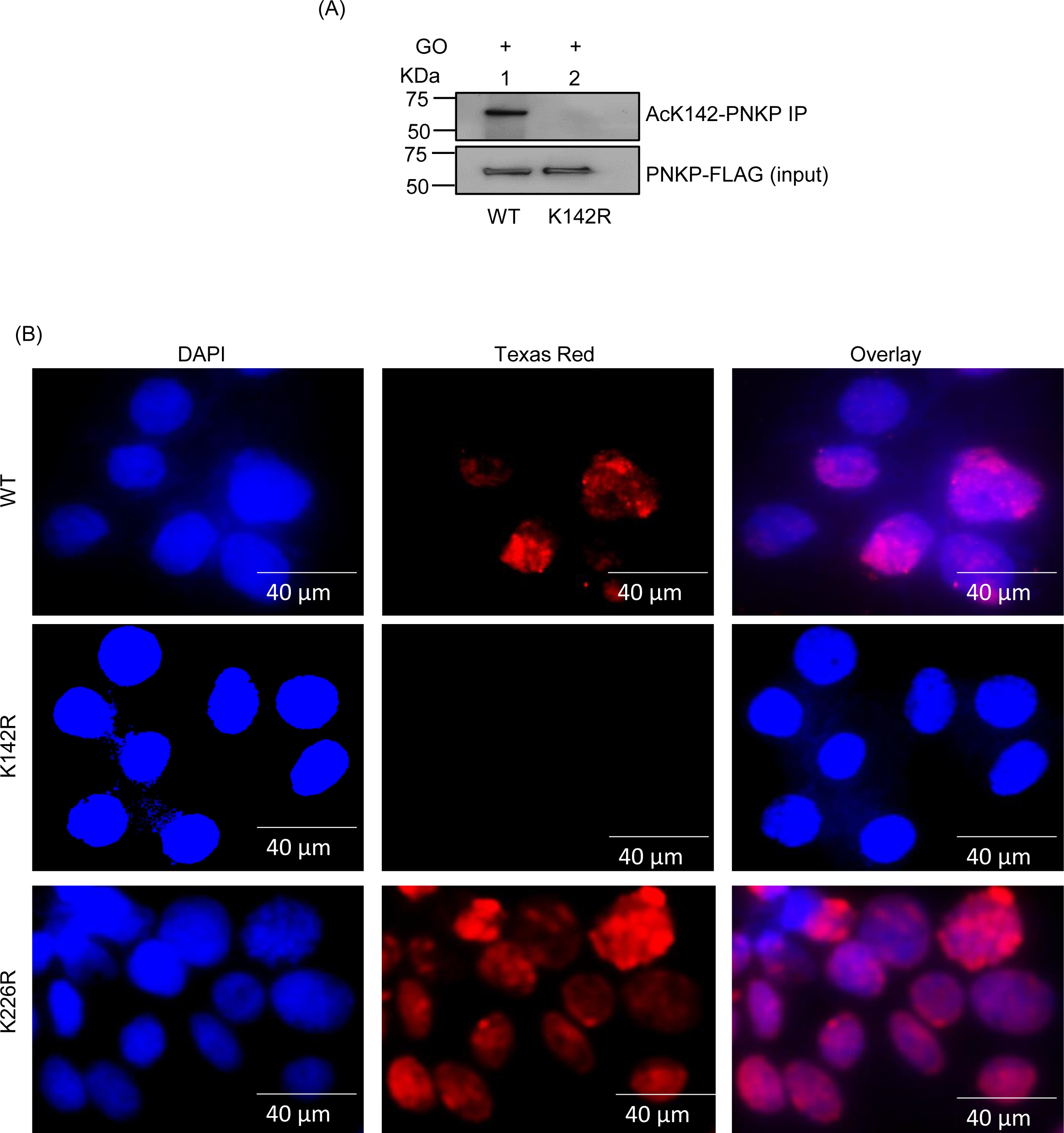
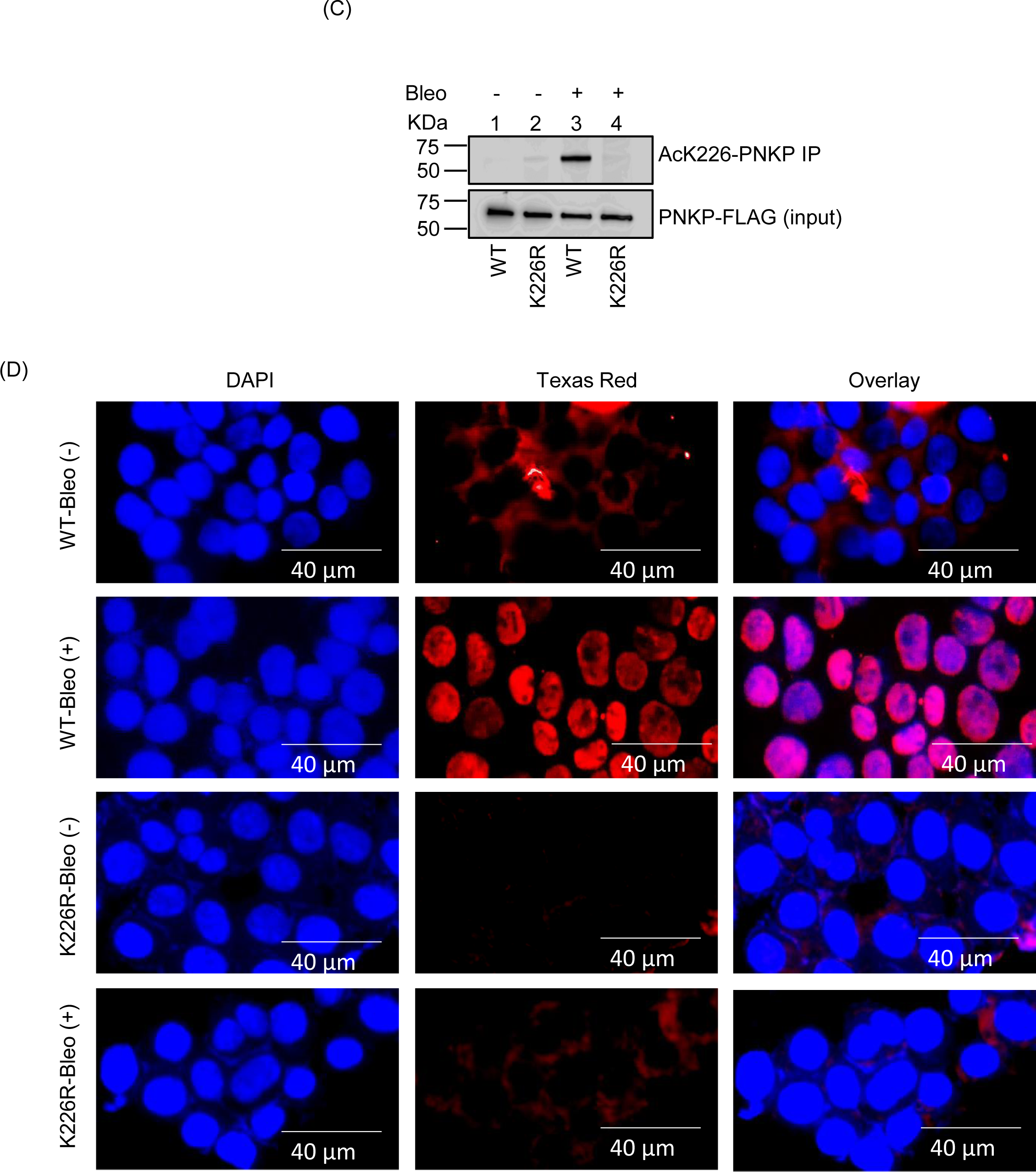
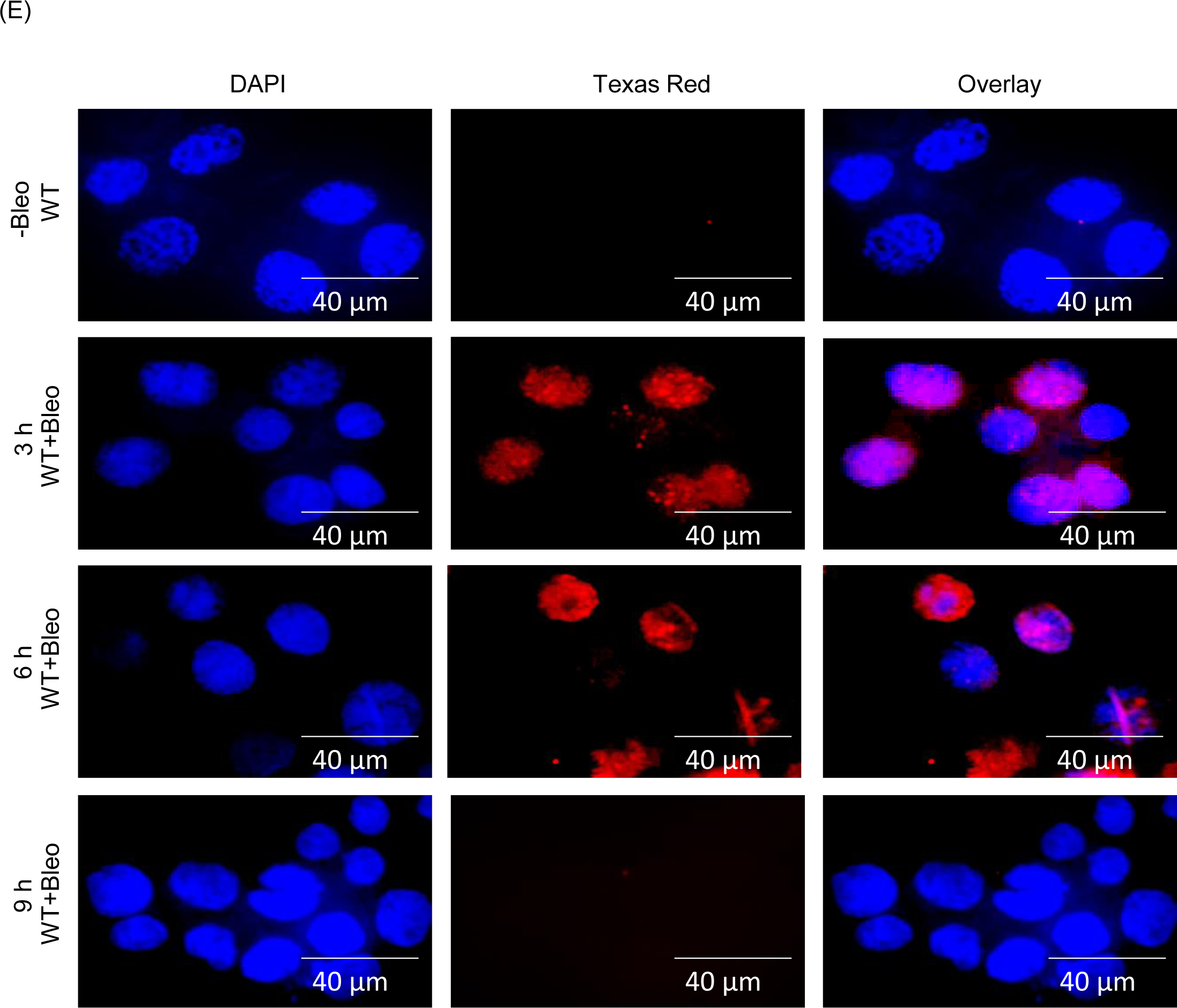
Validation of K142 and K226 acetylation sites using custom-generated anti-Ac (K142 or K226) specific antibodies (Ab) in stable cell lines. (A) PNKP was IP’d using the anti-AcK142 Ab from the chromatin fraction of GO-treated WT (lane 1) and K142R mutant (lane 2) PNKP expressing cells and probed with the anti-FLAG Ab (upper panel). Lower panel: Input shows similar ectopic expression of PNKP-FLAG in the cell lines, detected with the anti-FLAG Ab. (B) Indirect immunofluorescence was conducted to detect nuclear localization of AcK142 PNKP in GO-treated WT vs. mutant (K142R and K226R) cell lines. Nuclei were counterstained with DAPI. Images were taken at 40 µm area. (C) PNKP was IP’d using the anti-AcK226 Ab from the chromatin fraction of mock (lanes 1, 2) or Bleo-treated (lanes 3, 4) WT (lanes 1, 3) and K226R mutant (lanes 2, 4) PNKP expressing cells and probed with anti-FLAG Ab (Upper Panel). Lower panel shows the inputs, indicating similar ectopic expression of PNKP-FLAG in the cell lines detected with anti-FLAG Ab. (D) Mock- or Bleo-treated WT and mutant K226R PNKP expressing cells were stained with an anti-AcK226 Ab. Nuclei were counterstained with DAPI. (E) Indirect immunofluorescence was performed to assess the status of AcK226 PNKP in mock vs. Bleo-treated WT-PNKP expressing cells at 3 h, 6 h and 9 h post-Bleo treatment. All the immunofluorescence experiments were performed following depletion of endogenous PNKP by 3’-UTR specific siRNA.

Next, we performed a DSB-repair time course experiment following Bleo treatment in WT-PNKP expressing stable HEK293 cells (17–19) and found that acetylation at K226 was significantly reduced at 9 h post-Bleo treatment (**Fig. 1E, bottom panel)**, indicating a reversal of K226 acetylation with the progression of DSB repair. These data not only confirmed DSB-induced acetylation but also highlighted the reversible nature of the K226 acetylation event in PNKP.

### p300 acetylates PNKP at lysine 142 and CBP acetylates at lysine 226

p300 and CBP are two major histone acetyltransferases (HATs) responsible for the acetylation of the vast majority of proteins *in vivo* (35–37). To explore any potential role of these HATs in the acetylation of PNKP in mammalian cells, we individually depleted the endogenous p300 **(Supplementary** Fig. 3A) and CBP **(Supplementary** Fig. 3B) in HEK293 cells using commercially available siRNAs. We then performed indirect immunofluorescence using acetylation-specific Abs to assess the acetylation status at distinct residues. Immunofluorescence analyses with anti-Ac-K142 Ab revealed the presence of AcK142 PNKP in control HEK293 cells either mock-treated or treated with GO (**Fig. 2A, top two panels, respectively)**. AcK142 PNKP was also detected in the nucleus of CBP-depleted HEK293 cells but not in p300-depleted cells (**Fig. 2A, bottom two panels, respectively)**. This clearly indicated that the acetylation of PNKP at lysine 142 residue is mediated by p300, but not by CBP. Similarly, immunofluorescence analyses with anti-AcK226 Ab showed the presence of AcK226 PNKP in Bleo-treated control cells (**Fig. 2B, 2_nd_ panel from the top)** but not in mock-treated control cells (**Fig. 2B, top panel)**, consistent with our earlier observation of DSB-induced K226 acetylation. Most importantly, no such acetylation of PNKP was observed in Bleo-treated CBP depleted cells (**Fig. 2B, 3rd panel from the top)**, unlike in p300 depleted/Bleo-treated cells (**Fig. 2B, bottom panel)**. This finding confirmed that the DSB-induced acetylation of PNKP at K226 is mediated by CBP but not by p300. These experiments unveil the role of p300 and CBP in two distinct site-specific acetylation events of PNKP, viz., K142 and K226, respectively.

**Figure 2.**
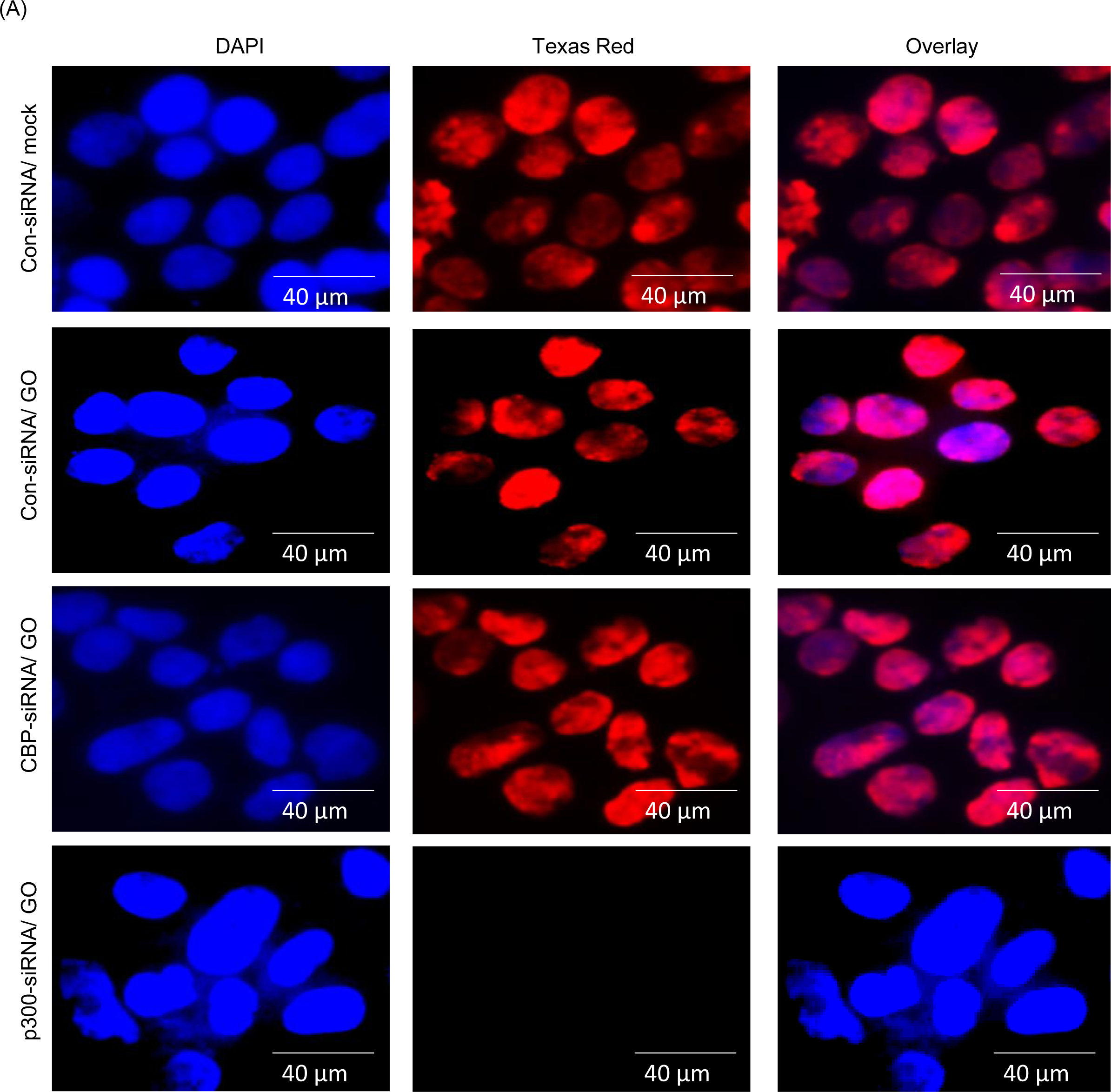
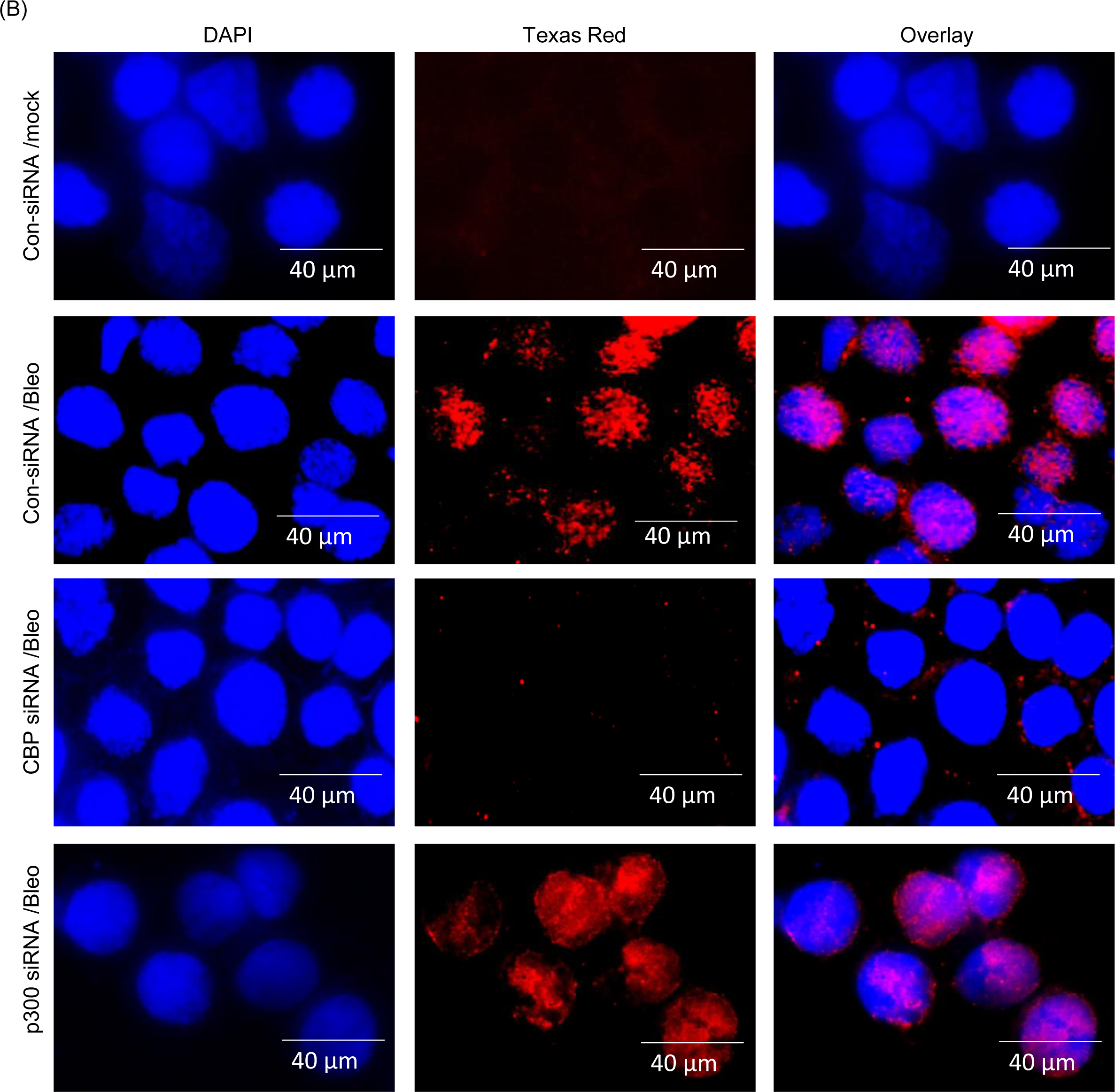
Identification of Histone Acetyl Transferases (HATs) responsible for acetylation of PNKP. (A) Detection of K142 specific PNKP acetylation in control (Con) siRNA vs. CBP/p300 siRNA transfected and mock/GO-treated HEK293 cells by indirect immunofluorescence using anti-AcK142 PNKP Ab. The three separate panels show nuclear DAPI staining, Texas red Ab staining, and the overlay. (B) K226-specific PNKP acetylation in control siRNA vs. CBP/p300 siRNA transfected ± Bleo-treated HEK293 cells was determined by fluorescence microscopic imaging using anti-AcK226 PNKP Ab.

### Acetylation of PNKP at K226 caused modest activation of its 3’-phosphatase but not kinase activity

PNKP can convert the blocked 5’-OH and 3’-P DNA termini to canonical 5’-P and 3’-OH DNA termini, respectively, enabling the gap-filling and ligation of DNA ends (6–10). Thus, we explored the effect of these acetylation events on PNKP’s 3’-phosphatase and 5’-kinase activities using affinity-purified FLAG-PNKP from control vs. Bleo-treated stable cell lines (WT, K142R, K226R and K142R/K226R). Bleo-induced DSB generation was confirmed by the expression of DSB marker, γH2AX in the chromatin fraction of WT **(lanes 1-2)** and mutant **(lanes 3-4,** K142R**; lanes 5-6,** K226R; and **lanes 7-8,** K142R/K226R) FLAG-PNKP expressing cells (**Fig. 3A, upper panel)**. PNKP was IP’d from the same extract with anti-PNKP Ab and immunoblotted with anti-FLAG Ab to confirm the similar level of FLAG-PNKP expression in these cells following mock/Bleo-treatment (**Fig. 3B**). Assessment of 3’-phosphatase activity **(schematically shown in Fig. 3C**) revealed a modest activation of PNKP in WT **(lanes 3 vs. 2)** and K142R **(lanes 5 vs. 4)** cells (K226 acetylation proficient) following Bleo-treatment, whereas no significant change of activity was observed in Bleo-treated K226R **(lanes 7 vs. 6)** and K142R/K226R **(lanes 9 vs. 8)** PNKP expressing cells (**Fig. 3D**). This observation was further confirmed by analyzing the 3’-phosphatase activity of affinity-purified PNKP from WT vs. acetylation-mimic (K142Q, K226Q, K142Q/226Q) PNKP expressing cells that showed no difference in activity between them (**Fig. 3E**). Additionally, we assessed 5’-kinase activity using affinity-purified PNKP from WT vs. acetylation-deficient (K-R) mutant cells and also observed no difference in such activity among the cell lines ± Bleo (**Fig. 3F).** Overall, these experimental findings clearly indicated that acetylation at K226 induces a modest activation of PNKP’s 3’-phosphatase activity but does not affect its kinase activity.

**Figure 3.**
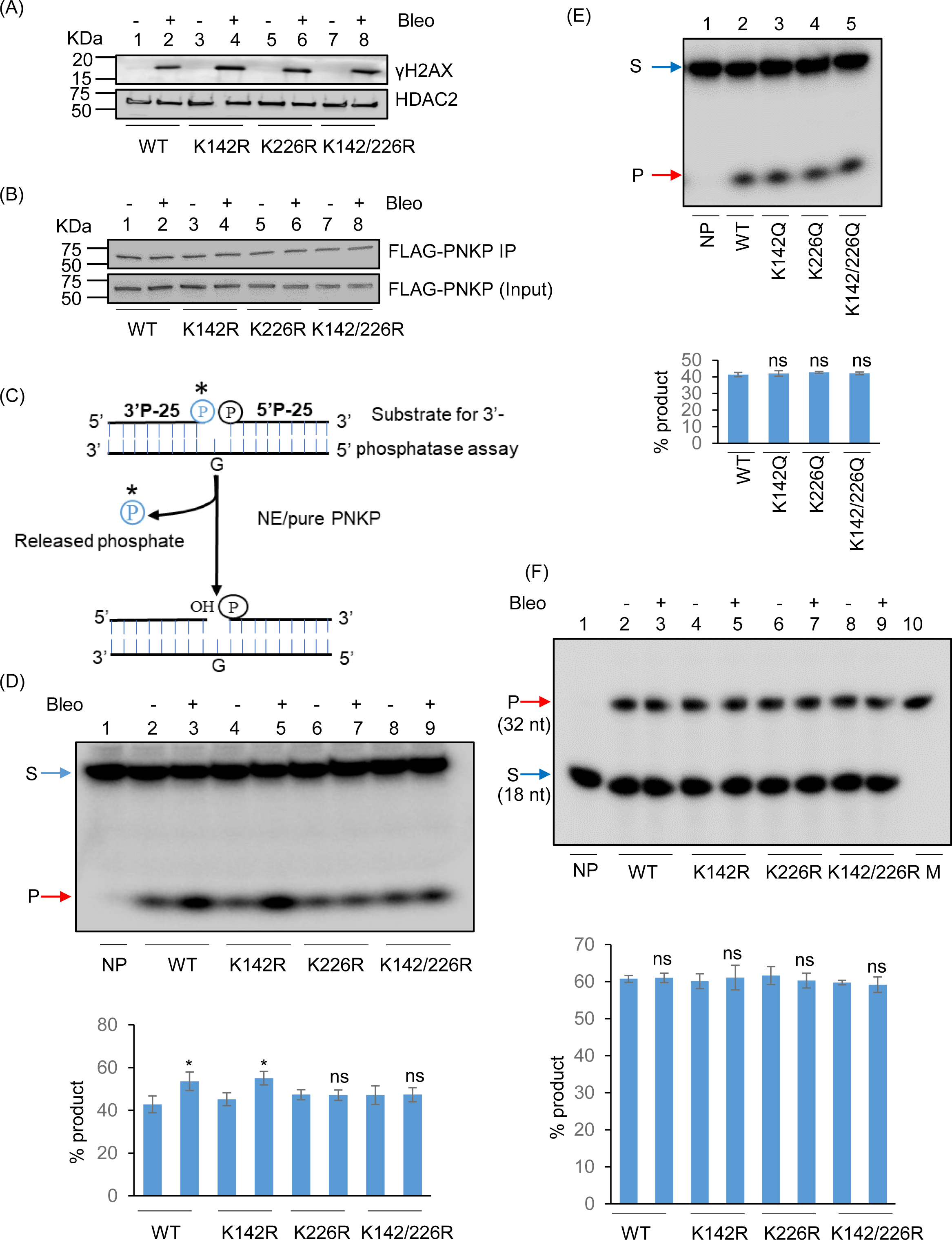
Assessment of the effect of acetylation on PNKP’s 3’-phosphatase and 5’-kinase activities. (A) The Western blot shows the expression of γH2AX (upper panel) in the chromatin fraction of WT (lanes 1, 2), K142R (lanes 3, 4), K226R (lanes 5, 6) and K142R/226R (lanes 7, 8) PNKP expressing cells post mock (-) or Bleo (+) treatment. HDAC2 is used as the loading control for the chromatin fraction (lower panel). (B) IP’d and purified FLAG-PNKP from WT and mutant PNKP expressing cells (± Bleo) was immunoblotted with anti-FLAG Ab (upper panel). 10 µg of the IP’d chromatin fraction from WT and K-R mutant PNKP expressing cells was used as input (lower panel). (C) A schematic representation of the PNKP 3’-phosphatase assay. (D) The 3′-phosphatase assay was performed with immuno-purified FLAG-PNKP from WT (lane 2-3), K142R (lane 4-5), K226R (lane 6-7) and K142R/226R (lane 8-9) PNKP-expressing cells post mock (-) or Bleo (+) treatment. Lane 1: No protein (NP), substrate only. Lower panel represents the quantitation of the products (% of released phosphate) (Error bars show ±SD of the mean; n=3, *P<0.05 between mock (-) and Bleo (+) treatment in WT and K142R; ns: non-significant, P>0.05). (E) The phosphatase assay was performed with immuno-purified FLAG-PNKP from WT (lane 2), K142Q (lane 3), K226Q (lane 4) and K142Q/226Q (lane 5). Lane 1: no protein (NP), substrate only. Lower panel shows the quantitation of the products (% of released phosphate) (Error bars show ±SD of the mean; n=3, ns: non-significant, P>0.05). (F) 5′-kinase assay was conducted with IP’d FLAG-PNKP from WT (lane 2-3), K142R (lane 4-5), K226R (lane 6-7) and K142R/226R (lane 8-9) post mock (-) or Bleo (+) treatment. Lane 10: radiolabeled 32 nt marker (M). Lane 1: No protein (NP), substrate only. The quantitation of the products (% phosphorylated product) is represented in the bar diagram (Error bars show ±SD of the mean; n=3, ns=non-significant, P>0.05). In each case of (D-F), S: Substrate and P: product.

### Acetylation at K142 plays a critical role in repairing SSBs in the transcribed genes

To investigate the role of K142 and K226 acetylated PNKP in SSB repair, WT, K-R (acetylation-deficient) and K-Q (acetylation-proficient/mimic) mutant PNKP expressing cells were treated with Glucose Oxidase (GO) to induce SSBs, and the strand-break accumulation was analyzed in representative transcribed (HPRT, Pol Beta and RNA polII) vs. non-transcribed (NANOG, OCT4 and MyH2) genes using long-amplicon qPCR (LA-qPCR) (17,18), after allowing cells to recover for 4 h following DNA damage induction. To specifically assess the role of ectopically expressed PNKP in DNA SSB repair, we performed prior depletion of endogenous PNKP in these cells by 3’ UTR specific siRNA **(Supplementary** Fig. 4A**; supplementary Fig. 4B)**. It was found that efficient SSB repair of transcribed genes occurred in WT (**Fig. 4A, lanes 1-3, upper panel)** and K226R (**Fig. 4A, lanes 7-9, upper panel)** cells, but such repair was significantly abrogated in K142R (**Fig. 4A, lanes 4-6, upper panel)** and K142R/226R (**Fig. 4A, lanes 10-12, upper panel)** mutant PNKP expressing cells. Interestingly, SSB repair was not affected in non-transcribed genes (**Fig. 4A, lower panel)** despite similar SSB levels compared to transcribed genes (**Fig. 4A Upper and Lower Panels, lanes 2, 5, 8, 11**) after GO treatment and we observed almost complete repair in WT **(lanes 1-3)**, K142R **(lanes 4-6)**, K226R **(lanes 7-9)** and K142R/226R **(lanes 10-12)** cells. These findings are consistent with our earlier reports of PNKP’s involvement in the preferential repair of transcribed genes (17–19). Thus, we conclude that, the repair of SSBs in transcribed genes indeed relies mostly on K142 acetylation but not on K226 acetylation. This conclusion was further corroborated with our observation of complete repair of transcribed genes in acetylation-mimic (K-Q) PNKP-expressing cells compared to WT (**Fig. 4B**). Collectively, our data suggest that AcK142 PNKP plays a critical role in the SSB repair of transcribed regions.

**Figure 4.**
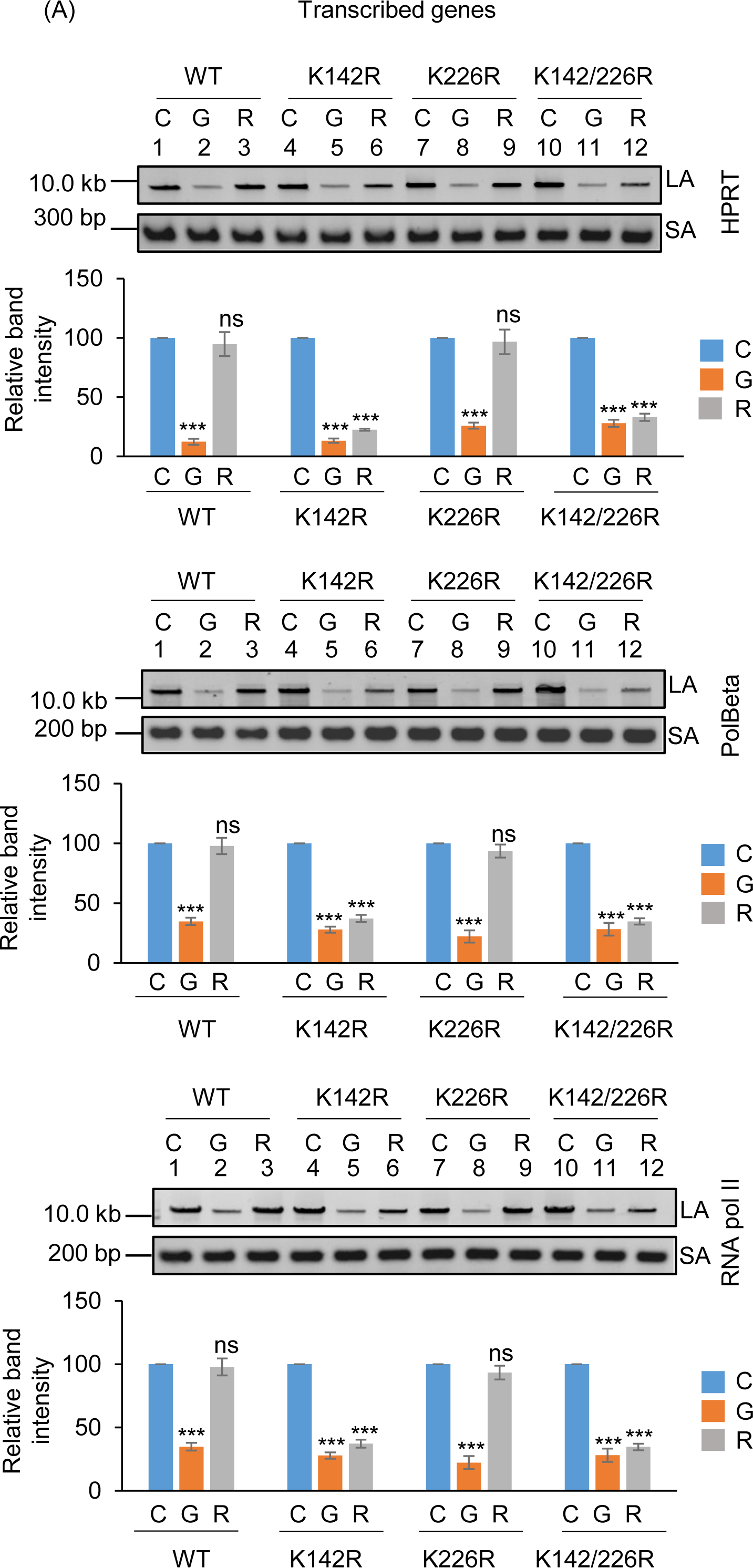
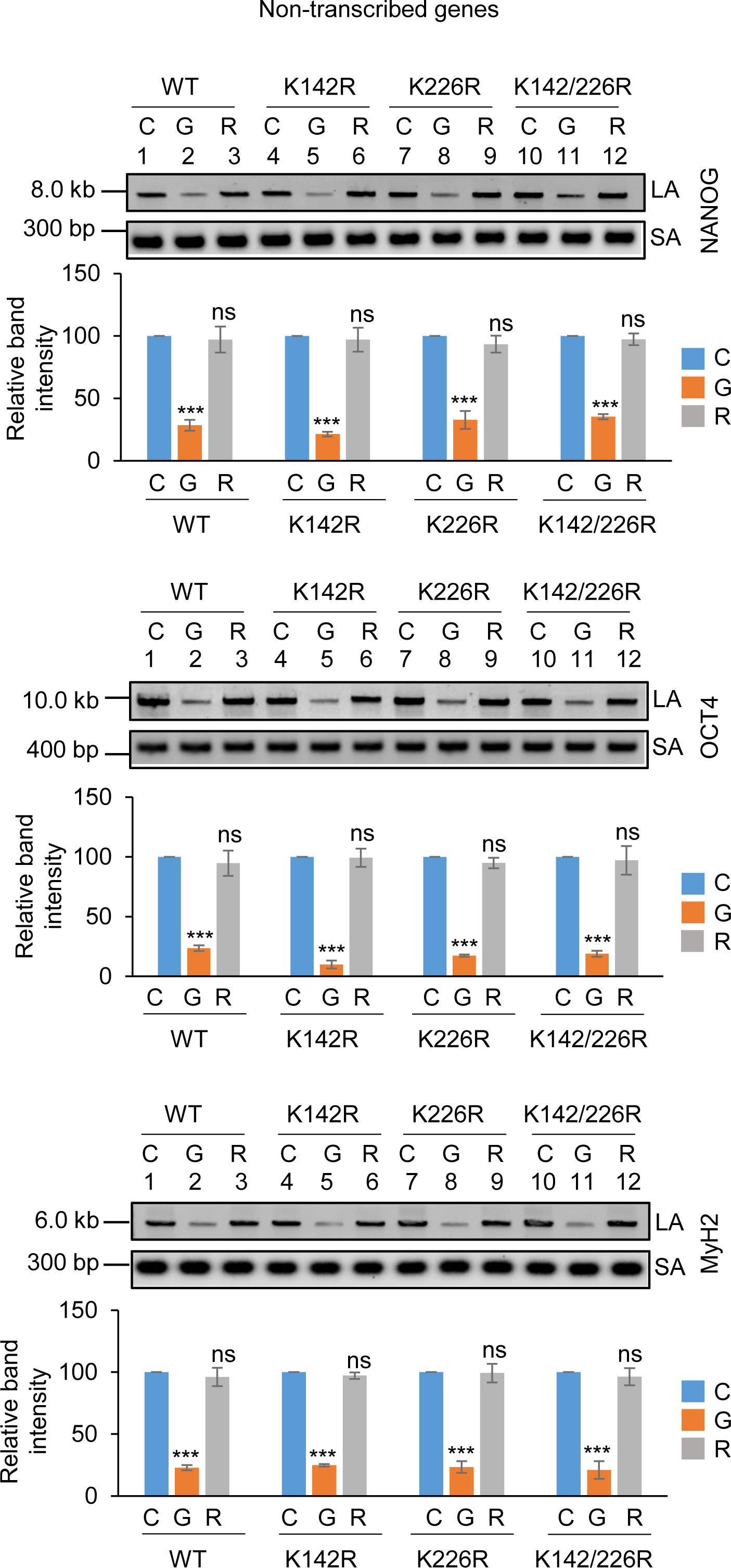
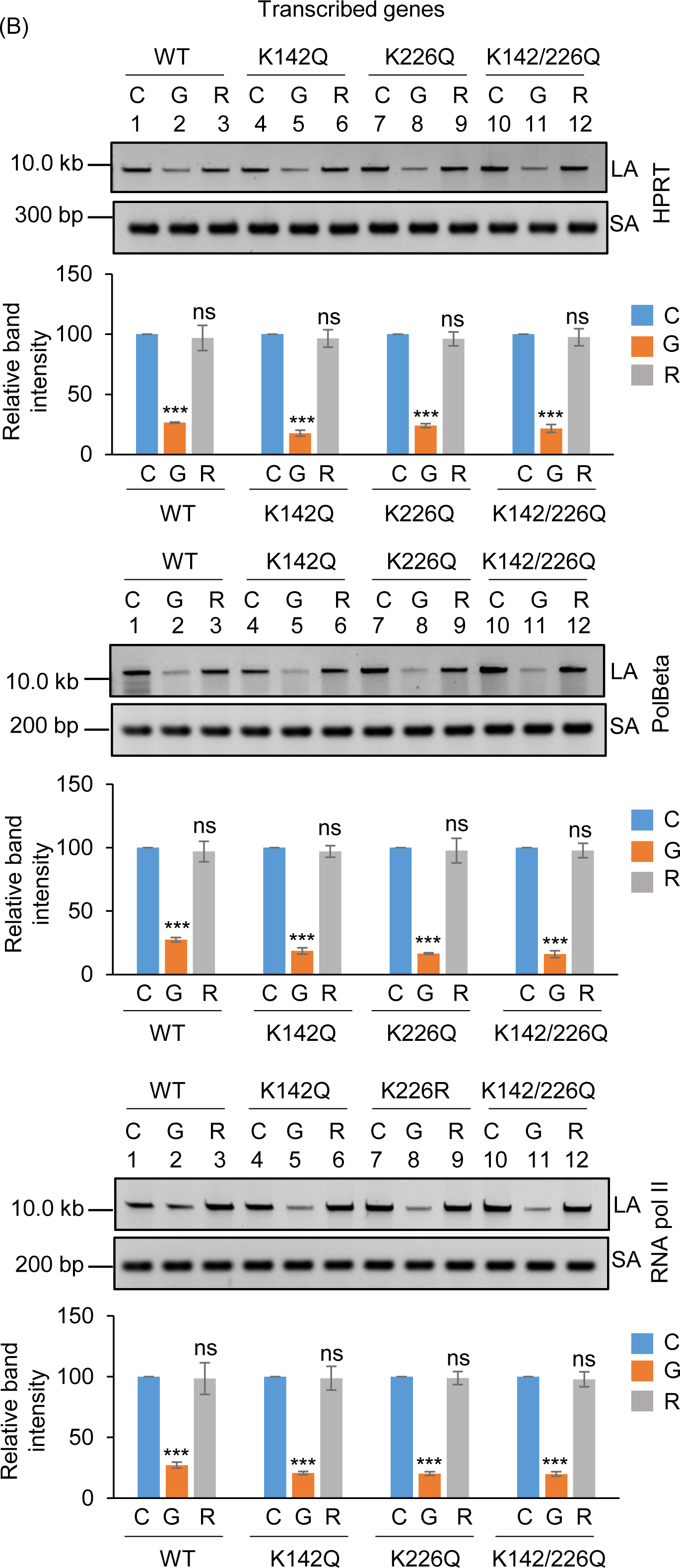
The role of lysine 142 acetylated PNKP on SSB repair. (A) (Upper panels) Representative agarose gel images showing each long amplicon (10–12 kb) and a short amplicon (∼200–300 bp) of the transcribed (HPRT, Pol Beta, and RNA polII) genes and non-transcribed (NANOG, OCT4 and MyH2) from the genomic DNA isolated from WT (lanes 1-3), K142R (lanes 4-6), K226R (lanes 7-9) and K142R/226R (lanes 10-12) PNKP expressing cells either mock (C), GO-treated (G), or at 4 h post-GO treatment (R). (Lower Panels) The bar diagrams represent the normalized (with short PCR amplicon) relative band intensity with the mock-treated control (C) set as 100 (Error bars show ±SD of the mean; n=3, ***P<0.005; ns=non-significant, P>0.05) for each cell line. (B) Similar LA-qPCR assay of the transcribed (HPRT, Pol Beta, and RNA polII) genes from the genomic DNA isolated from WT (lanes 1-3), K142Q (lanes 4-6), K226Q (lanes 7-9) and K142Q/226Q (lanes 10-12) PNKP expressing cells under similar treatment conditions as described in (A). All LA-qPCR experiments were performed following the depletion of endogenous PNKP in WT and mutant cells by 3’-UTR-specific siRNA.

### Preferential repair of DSBs in the transcribed genes requires acetylation at K226 of PNKP

To further understand the functional significance of the two distinct acetylated residues (K142 and K226) of PNKP in SSB vs. DSB repair, we assessed the DNA strand-break (SB) accumulation in WT and mutant (K-R and K-Q) PNKP expressing cells by gene-specific LA-qPCR after the cells were treated with Bleo (to induce DSBs predominantly) and allowed 12-16 h for repair. As in the previous case, we performed prior depletion of endogenous PNKP in these cells by 3’ UTR specific siRNA **(Supplementary** Figs. 5A**, C)**. We observed significant accumulation of DNA strand break in both transcribed (**Fig. 5A, upper panel)** and non-transcribed (**Fig. 5A, lower panel)** genes in all these cells after Bleo treatment. However, DNA SBs were almost completely repaired in WT **(lanes 3 vs. 1)** cells, whereas K142R **(lanes 6 vs. 4)**, K226R **(lanes 9 vs. 7)** and K142R/K226R **(lanes 12 vs. 10)** mutant PNKP expressing cells showed persistence of DNA SBs in the transcribed genes (**Fig. 5A, upper panel)**. In contrast, almost complete repair of DNA strand breaks was observed in non-transcribed genes in K142R **(lanes 6 vs. 4)**, K226R **(lanes 9 vs. 7)** and K142R/K226R **(lanes 12 vs. 10)** mutant cells (**Fig. 5A, lower panel)**. Our results thus suggest that acetylated (both K142 and K226 specific) PNKP preferentially repaired the transcribed genes. Since Bleo treatment can induce both SSBs and DSBs (38–40), K142R mutant expressing cells also showed persistence of DNA SBs due to impairment of AcK142-mediated SSB repair. However, K226R PNKP expressing cells accumulate a significant amount of DNA SBs, unlike what was observed following GO-mediated SSB induction and subsequent repair. It is therefore evident that acetylation at the K226 site is primarily involved in DSB repair. Consistently, we observed almost complete resolution of DSB marker, γH2AX in K142R PNKP expressing cells following repair, indicating efficient DSB repair in these cells. On the contrary, we observed persistent γH2AX in K226-acetylation deficient (K226R and K142R/K226R) cells, indicating impaired DSB repair **(Supplementary** Fig. 5B). Notably, K226Q (acetylation mimic) efficiently repairs the strand breaks in the transcribed genes, which is further substantiated by resolution of γH2AX (DSB marker) following repair (**Fig. 5B and Supplementary** Fig. 5D). These data further confirm the critical role of PNKP acetylation in the repair of DSBs in the transcribed genes.

**Figure 5.**
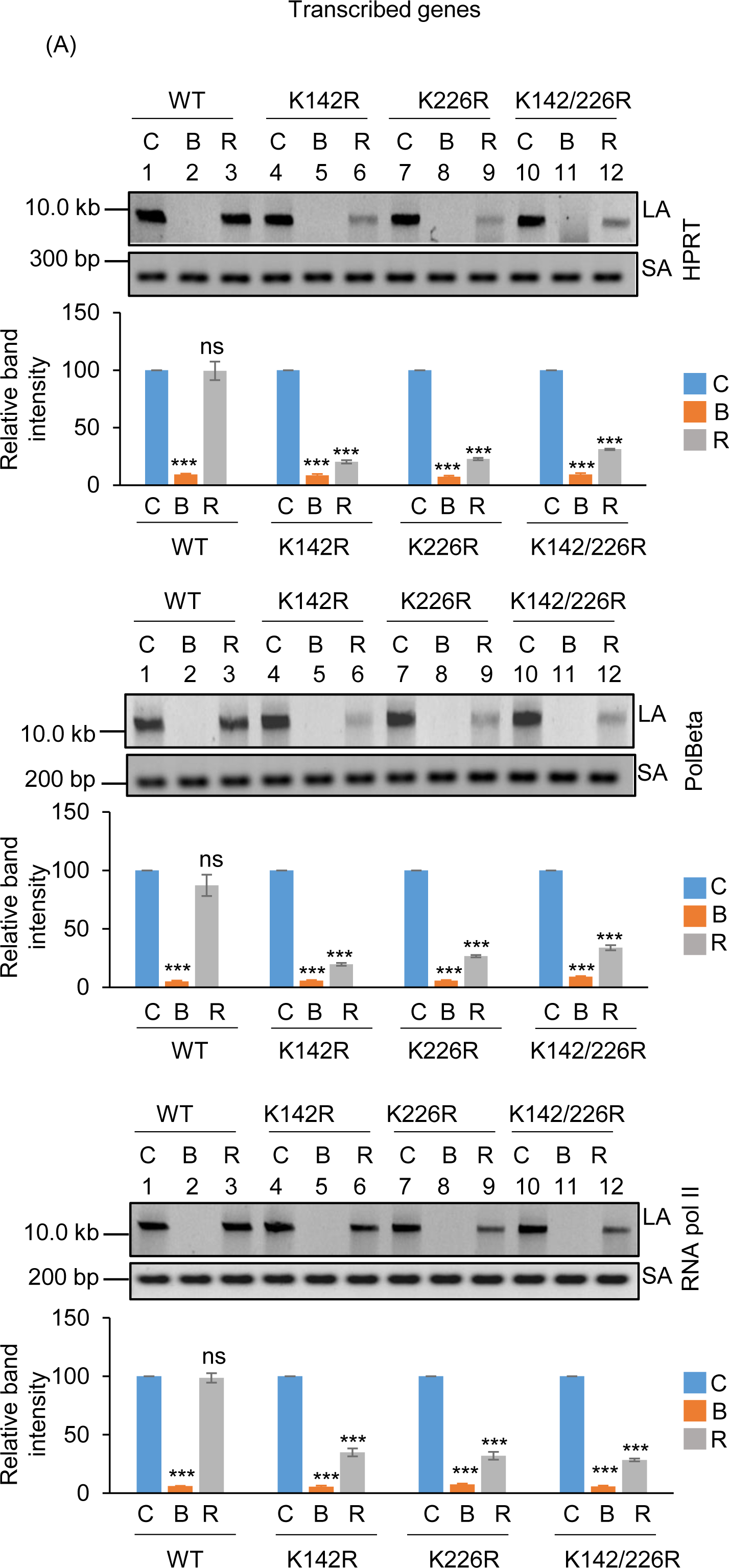
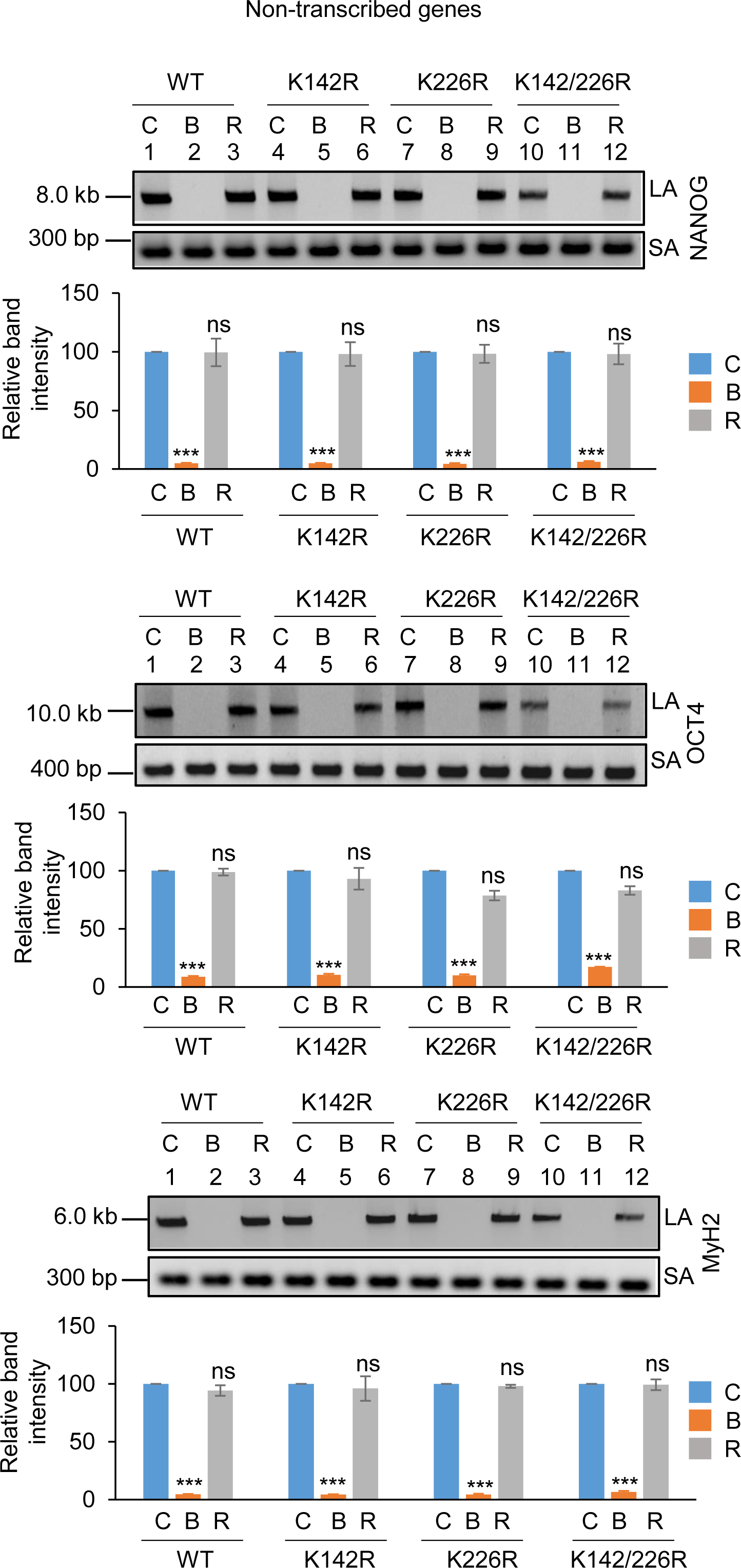
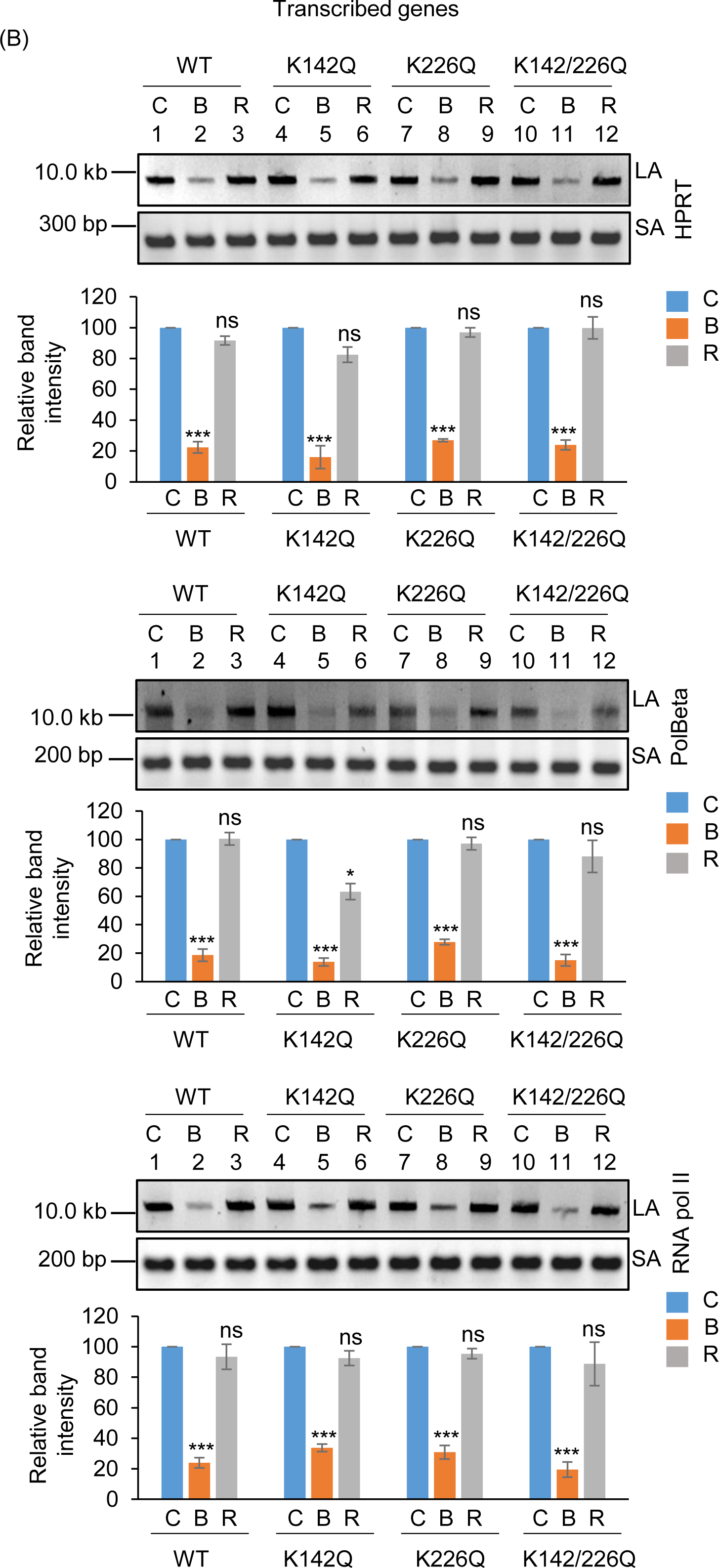
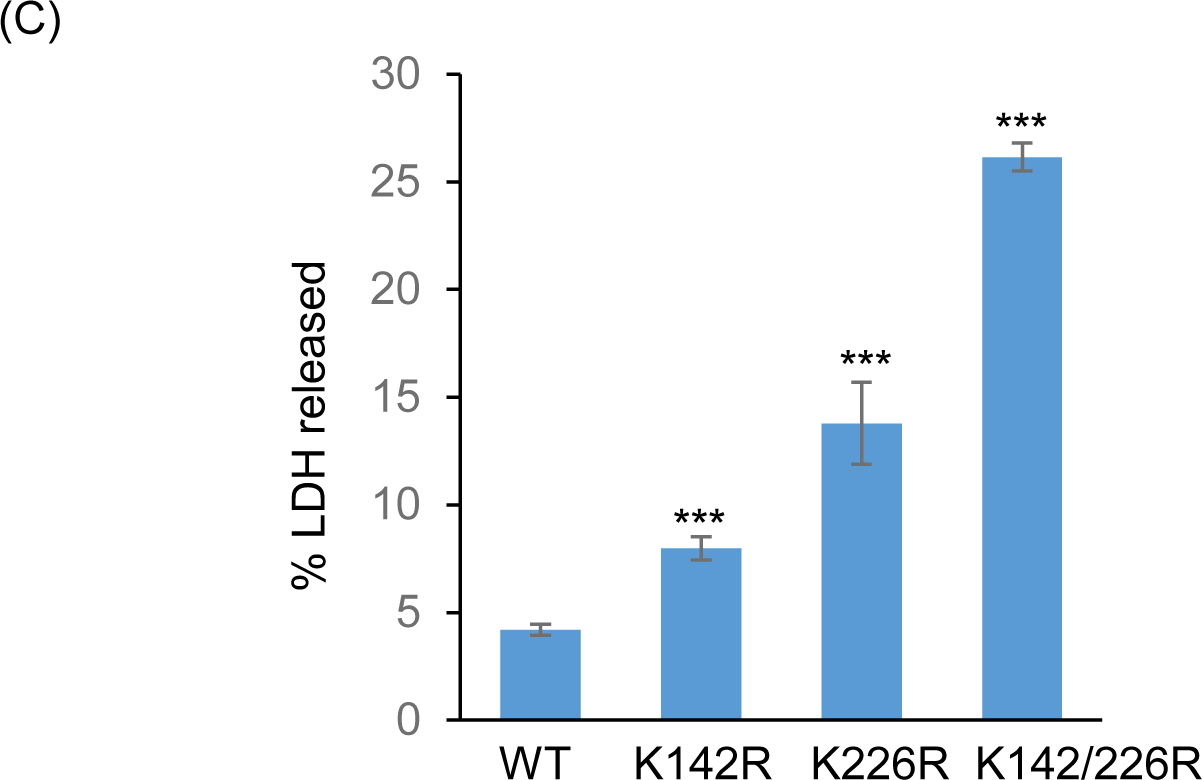
Assessment of the effect of lysine 226 specific PNKP acetylation on the repair of Bleo-induced DSBs by long amplicon qPCR. (Upper panels) Representative agarose gel images of the long amplicon (10–12 kb) and a short amplicon (∼200–300 bp) of the corresponding genes from WT (lanes 1-3), K142R (lanes 4-6), K226R (lanes 7-9) and K142R/226R (lanes 10-12) PNKP expressing cells either mock (C), Bleo-treated (B), or at 16 h (R) following Bleo-treatment. (Lower Panels) The bar diagrams represent the normalized (with short PCR amplicon) relative band intensity with the mock-treated control (C) set as 100 (Error bars show ±SD of the mean; n=3, ***P<0.005; ns: non-significant, P>0.05) for each cell line. (B) Similar LA-qPCR assay of the transcribed (HPRT, Pol Beta, and RNA polII) genes from the genomic DNA isolated from WT (lanes 1-3), K142Q (lanes 4-6), K226Q (lanes 7-9) and K142Q/226Q cells under similar treatment conditions as described in (A). (C) The bar graph represents the percent LDH release in K142R, K226R and K142R/226R mutant PNKP expressing cells compared to WT cells. Error bars show ±SD of the mean (n = 3, _∗∗∗_P < 0.005). All LA-qPCR/R-loop/LDH assays were performed following the depletion of endogenous PNKP in WT and mutant cells by 3’-UTR-specific siRNA.

Further, in view of abrogated DNA strand-break repair in mutant PNKP-expressing cells, we assessed the release of lactate dehydrogenase (LDH; a marker of cytotoxicity) following Bleo-treatment. Results indeed showed a significant increase of LDH release in mutant cells compared to WT (**Fig. 5C),** indicating significant toxicity in PNKP-mutant cells, consistent with enhanced DNA SB accumulation in these cells.

### K142 or K226 acetylated PNKP forms distinct pathway specific repair complexes

Based on our experimental findings so far, we hypothesize that K142 acetylation of PNKP plays a role in SSB repair whereas K226 acetylation is involved in DSB repair. In our earlier studies, we provided evidence of pre-formed, pathway-specific DNA repair complexes in mammalian cells (17–19). To investigate whether AcK142 PNKP and AcK226 PNKP associates to form a complex with SSBR and DSBR protein(s), respectively, we performed Co-IP from the chromatin fraction of WT-PNKP expressing stable cells (± GO or Bleo) using anti-AcK142 or anti-AcK226 Abs. Results indeed showed the presence of RNAP II, along with SSB repair proteins DNA Lig III and XRCC1 in the AcK142 PNKP immunocomplex, whereas DSB repair proteins, XRCC4 and DNA Lig IV were absent in this immunocomplex (**Fig. 6A**). On the contrary, we observed RNAP II, DNA Lig IV, Ku70 and XRCC4, but not the SSB repair proteins (DNA Lig III and XRCC1) in the AcK226 PNKP immunocomplex following Bleo-treatment (**Fig. 6B**). Importantly, we could specifically detect p300 and CBP in the K142 and K226 immunocomplexes, respectively, consistent with our immunofluorescence data showing their specific roles in acetylation of distinct lysines. These data further demonstrated the distinct role of PNKP acetylation at K142 and K226 residues on SSB and DSB repair pathways, respectively.

**Figure 6.**
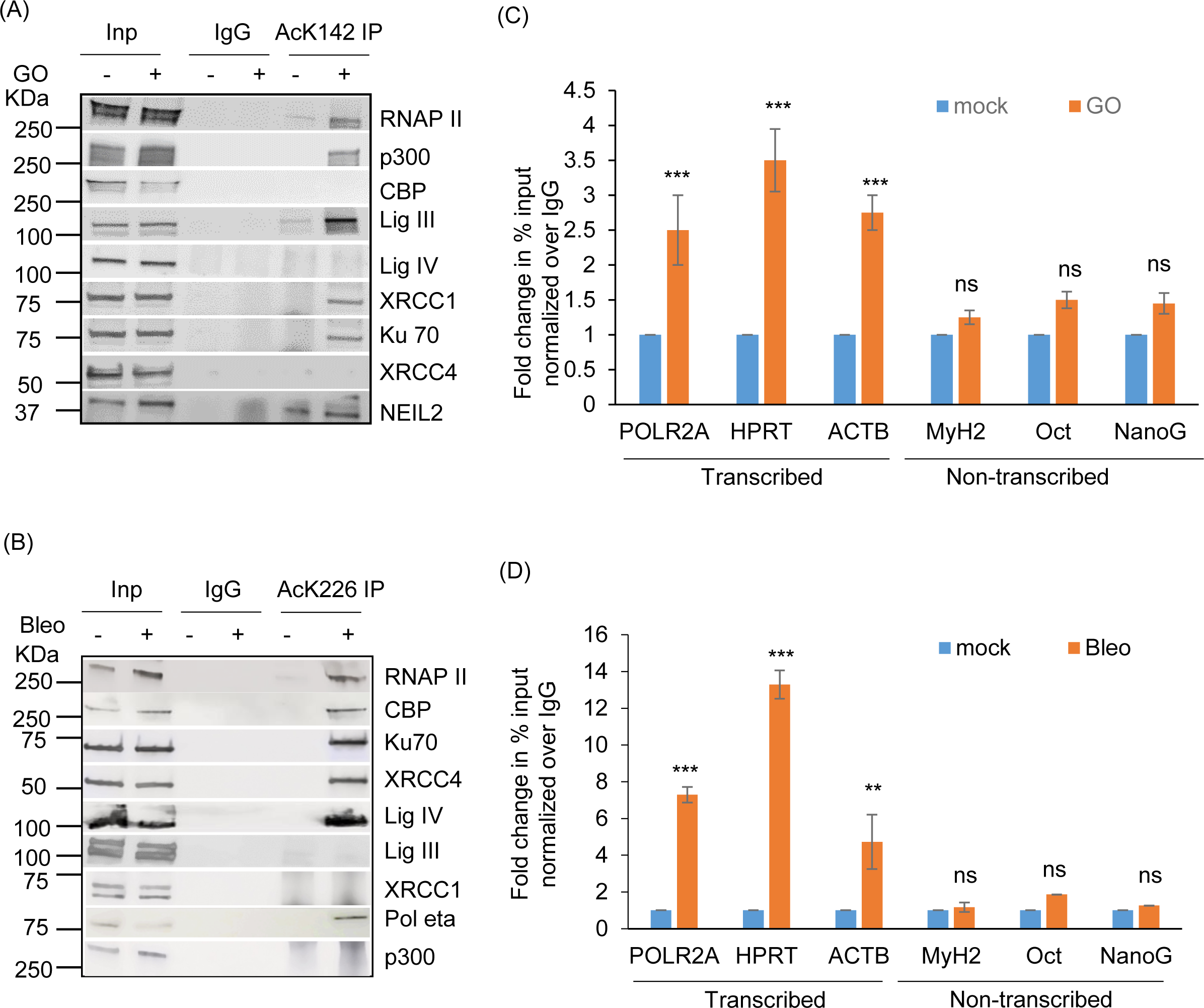
Characterization of AcK142 and AcK226 PNKP immunocomplexes and their association to a specific genomic region. Benzonase-treated chromatin fraction from the WT-PNKP-FLAG cells following mock (-)/GO(+) or mock(-)/Bleo(+) treatment was IP’d individually with (A) anti-AcK142 Ab or (B) anti-AcK226 Ab, respectively along with control IgG (rabbit) Ab and probed with respective Abs as indicated to the right of the panels. The binding to the exonic regions of transcribed (POLR2A, HPRT, Actin B) vs. the non-transcribed (MyH2, Oct3, NanoG) genes were quantified by quantitative PCR (qPCR) from IP’d DNA from PNKP-FLAG stable cells ± GO or ± Bleo treatment using AcK142 (C) or AcK226 (D) Abs, respectively. The data are represented as fold enrichment of % input over IgG with mock-treated samples considered as unity. Error bars represent ±SD of the mean (n = 3). ***P < 0.005, **P < 0.01 represent statistical significance between mock and GO/Bleo treatment for each gene (ns=nonsignificant, P > 0.05).

### Preferential association of acetylated PNKP with the transcribed regions

Since the Ac-PNKP preferentially repaired the transcribed genes, we examined the genomic association of AcK142 or AcK226 PNKP by performing ChIP on soluble chromatin preparations of WT-PNKP expressing stable cells (± GO or Bleo, respectively) using the AcK142 or AcK226 PNKP Abs. We observed the preferential recruitment of both AcK142 and AcK226 PNKP in the coding region of transcribed genes (POLR2A, HPRT and ACTB) but not in the non-transcribed genes (MyH2, Oct and NanoG) (**Fig. 6C-D**). From these and earlier findings, we conclude that in the event of SSB and DSB, AcK142 and AcK226 PNKP individually interact and form complexes with SSB and DSB repair proteins, respectively, and are preferentially recruited to the transcribed regions to facilitate repair.

### PNKP acetylation at K226 is impaired in Huntington’s Disease (HD) mouse-derived striatal neuronal cells (Q111)

Huntington’s disease (HD), a neurodegenerative disorder, is caused by aberrant expansion of polyQ repeat in the N-terminus of the Huntingtin (HTT) protein (24). Several earlier studies, including ours, have shown the specific degradation of CBP in HD (both cell culture and mouse models) (24,41–43). Therefore, we further investigated PNKP acetylation pattern in WT vs. HD mouse-derived striatal neuronal cells with normal and expanded polyQ repeats (Q7 and Q111, respectively) (44). Indirect immunofluorescence analysis using anti-AcK142 Ab in mock/GO-treated Q7 and Q111 cells showed the presence of acetylated PNKP in both Q7 (**Fig. 7A, top panel and 2_nd_ panel from the top)** and Q111 (**Fig. 7B, top panel and 2_nd_ panel from the top)** cells. However, when using anti-AcK226 antibody in mock vs. Bleo-treated cells, we observed the presence of acetylated PNKP only in Bleo-treated Q7 (**Fig. 7A, bottom panel**) cells. As expected, we could not detect any AcK226 PNKP in mock-treated Q7 cells (**Fig. 7A, 3_rd_ panel from the top)**. More interestingly, Bleo-treated Q111 (**Fig. 7B, bottom panel)** cells did not show acetylation of K226 residue, unlike Q7 cells, indicating impairment of such acetylation event in Q111 cells. We further assessed the levels of CBP, p300, γH2AX and PNKP in the chromatin fraction of Q7 and Q111 cells by immunoblotting using specific Abs. We observed a comparable level of p300 and PNKP (**Fig. 7C**) in both Q7 and Q111 cells but a significantly reduced level of CBP (**Fig. 7C**) in Q111 compared to Q7 cells. These data are consistent with our previous finding in the HD mouse and patient-derived iPSCs (24). Our results further corroborated the immunofluorescence data showing the presence of AcK142 PNKP in both Q7 and Q111 cells (as the p300 level is comparable) and impairment of Bleo-induced acetylation of K226 residue of PNKP in Q111 cells due to CBP degradation. Finally, we assessed the DNA damage accumulation in Q7 vs. Q111 cells in three representative transcribed genes (POLB, TUBB and NeuroD) and three non-transcribed genes (MyH4, MyH6 and MyoD) by LA-qPCR. The results showed a progressive and significant DNA strand break accumulation in transcribed genes in Q111 cells compared to Q7 cells (**Fig. 7D, E**). Consistently, a significantly elevated level of γH2AX in Q111 compared to Q7 cells (**Fig. 7C**) confirmed the DSB accumulation in Q111 cells. Collectively, these data are consistent with our earlier report of the accumulation of DNA SBs in transcribed genes in HD and further provided a mechanistic basis for DNA damage accumulation in the genomes of HD patients due to a lack of CBP-mediated K226 acetylation of PNKP.

**Figure 7.**
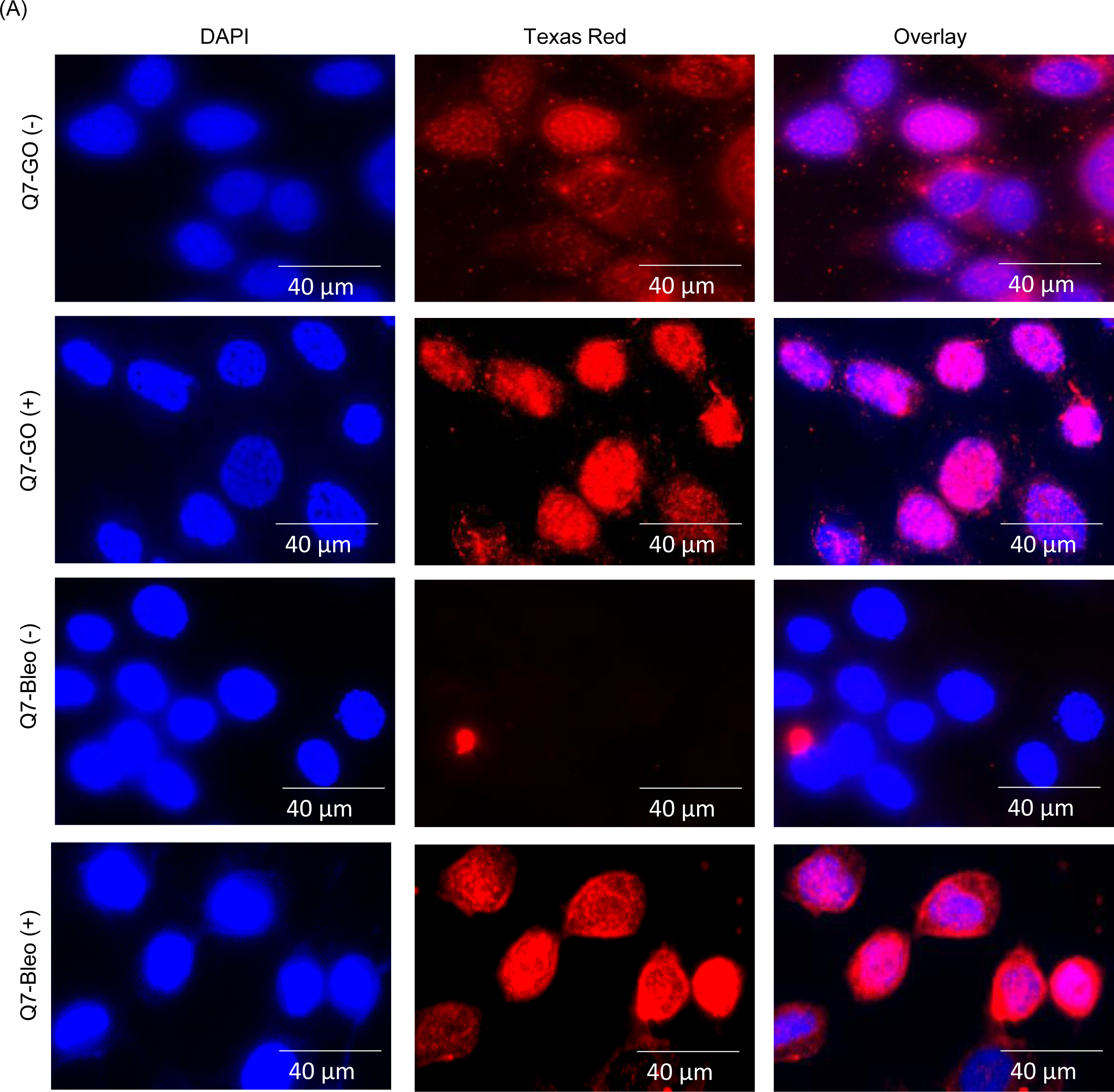
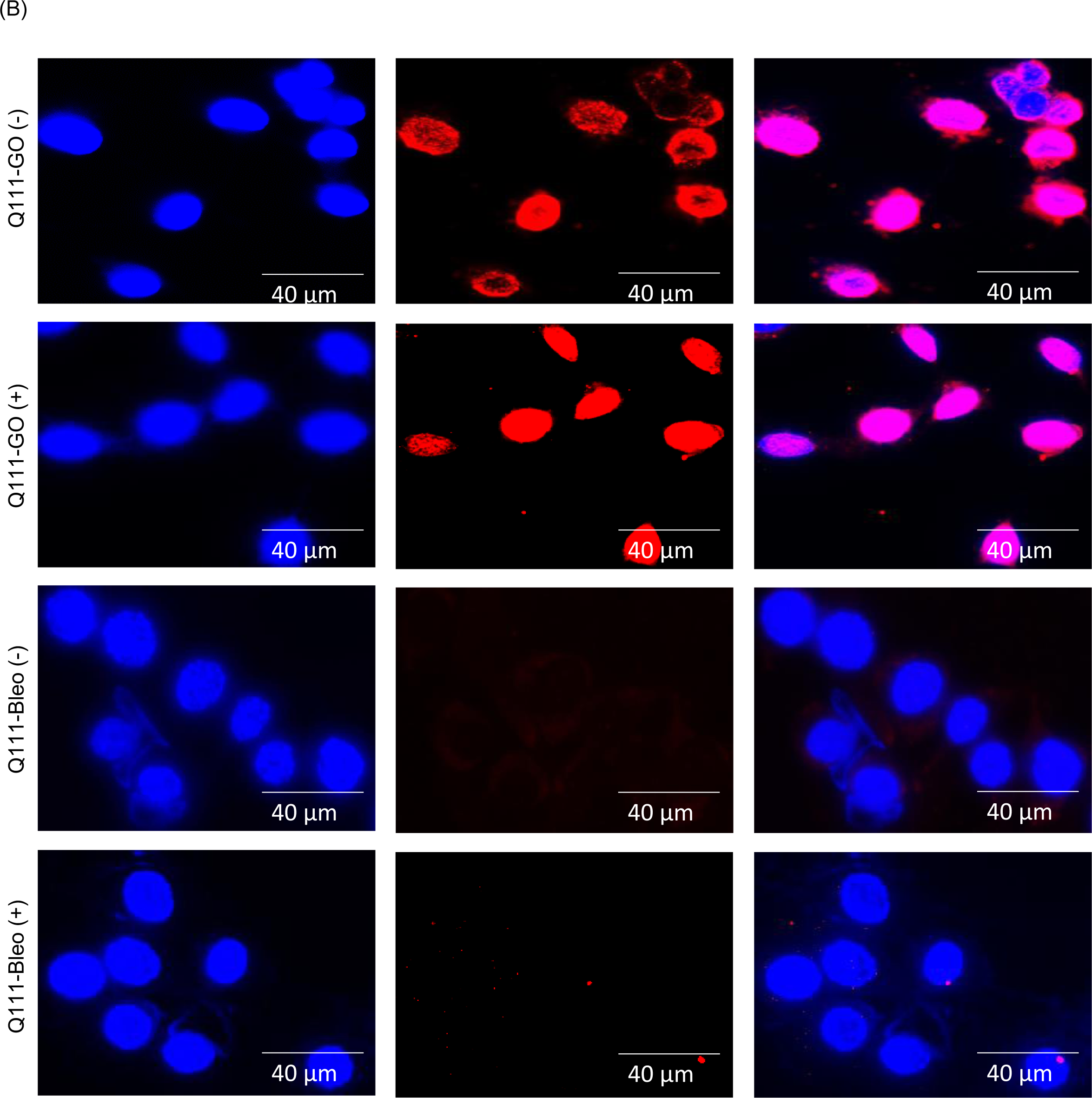
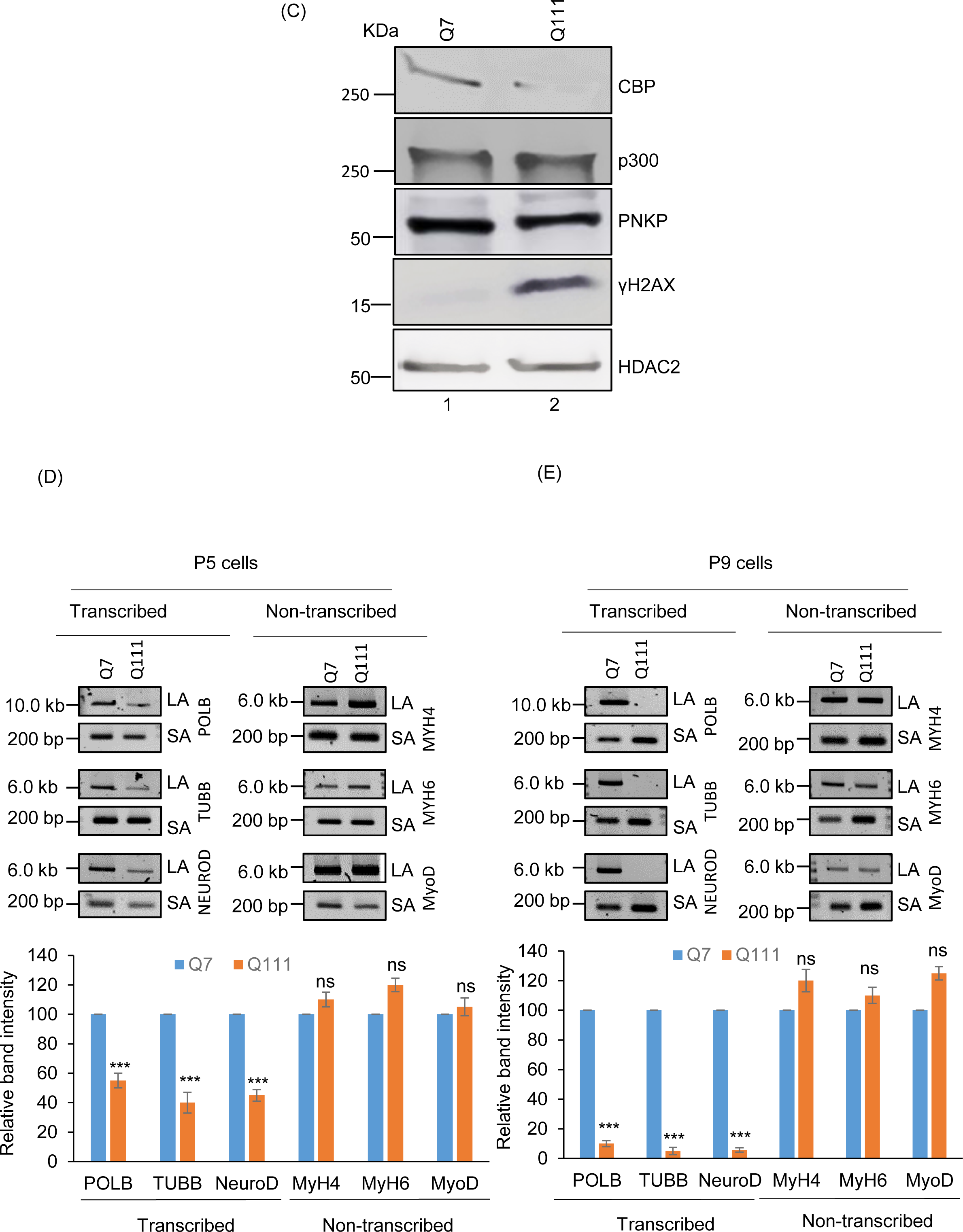
Assessment of K142 and K226 specific PNKP acetylation and estimation of DNA strand breaks in WT (Q7) vs. Huntington’s Disease (HD)-derived (Q111) striatal neuronal cell. (A) Detection of K142 specific acetylated PNKP (± GO) and K226 specific acetylated PNKP (± Bleo) in Q7 cells. Nuclear DAPI staining, Texas red Ab staining, and the overlay are shown side by side in three separate panels. (B) Similar microscopic imaging in Q111 cells ±GO/Bleo using anti-AcK142 and AcK226 Abs. (C) Western blots to assess the levels of proteins (as indicated on the right) in the chromatin fraction of Q7 vs. Q111 cells. HDAC2: loading control. (D-E) LA-qPCR for determining the DNA strand break accumulation in transcribed (POLB, TUBB and NEUROD) vs. non-transcribed (MYH4, MYH6 and MyoD) genes in Q7 vs. Q111 cells. (Upper panels): Representative agarose gel images of the Long (LA) vs. Short (SA) amplicons of DNA from Q7 and Q111 cells at passage 5 (P5) (D) and 9 (P9) (E). (Lower panels): Bar diagrams show the normalized (with short PCR amplicon) relative band intensity with the Q7 control set as 100 (Error bars represent ±SD of the mean; n=3, ***P<0.005, ns: non-significant, P>0.05).

## Discussion

The acetylation of protein is one of the most dominant post-translational modifications (PTMs) in eukaryotes and has a profound effect on the functional properties of multiple proteins that ultimately impact cellular physiology (45,46). Several human DNA replication and repair enzymes, involved in major DNA repair pathways, including mismatch repair (MMR), BER, nucleotide excision repair (NER), homologous recombination (HR), and NHEJ have been shown to be acetylated (5,11,13,28,47,48). Moreover, reversible acetylation of these proteins has been shown to play a key role in modulating DNA binding, transcriptional activation, protein stability and protein-protein interactions (49–51). Mitra’s group has previously reported the acetylation of several DNA repair enzymes involved in the BER pathway, such as NEIL1, NEIL2, OGG1 and APE1 (49–52). These acetylation events have diverse effects on the repair activity of the acetylated proteins. For instance, oxidative stress-induced acetylation significantly increases the activity of OGG1 in the presence of AP-endonuclease by decreasing its affinity for the product abasic (AP) site (52). Acetylation of NEIL2 inactivates its glycosylase/AP lyase activity (50), whereas acetylation of NEIL1 stabilizes the formation of chromatin-bound repair complexes that protect cells from oxidative stress (49). Acetylated NEIL1 preferentially associates with euchromatin and maintains transcribed genome integrity (53). NEIL2 is also involved in TC-BER (14,15); however, whether acetylation or any other modification plays any role in the involvement of NEIL2 in such repair pathway warrants further investigation.

The NEIL-initiated TC-BER pathway requires PNKP to process the 3’-P termini, one of the most abundant blocked termini at the site of DNA strand breaks. It has been shown that ionizing radiation (IR)-induced phosphorylation of PNKP, mediated by DNA-PKcs and ATM at selective serine residues, increases its DNA binding and phosphatase/kinase activities (21). Such modifications are important for the recruitment of PNKP to DNA damage sites but may not be the determining factor for the pathway choice of PNKP. We postulated that other types of PTMs could effectively modulate PNKP’s role in TC-BER/SSBR vs. TC-NHEJ. Here we report the identification of two acetylation sites (K142 and K226) in PNKP and demonstrate that acetylation at K142 occurs constitutively, and at K226 only after Bleo-induced DSB formation. Despite the acetylation deficient (K142R and K226R) as well as acetylation mimic (K142Q and K226Q) mutants having comparable enzymatic activity with that of WT PNKP, K142R and K226R expressing cells showed substantially higher level of DNA strand break accumulation in the transcribed genes only. However, acetylation mimic K142Q and K226Q mutants can efficiently repair transcribed genes. Moreover, DNA ChIP analysis showed a preferential association of the acetylated PNKP with the transcribed genome, indicating that acetylation of PNKP plays an important role in transcribed genome repair. These results are consistent with our earlier reports demonstrating PNKP’s preferential role in the repair of SSBs and DSBs in the transcribed genes (14,15,17,18).

Two major acetyltransferases, p300 and the transcriptional co-activator, CREB binding protein (CBP) are highly homologous (54–56) and both proteins are indispensable for development (57). While there is evidence of tissue-specific non-redundancy, in most cases CBP and p300 are considered to be functionally similar. We and others previously reported that p300 acetylates most of the DNA repair proteins involved in the BER process, such as FEN1, Polβ, TDG, NEIL1, NEIL2, APE1 and PCNA and modulate their activity, suggesting that acetylation of repair proteins contributes to the regulation of the BER process (49–51,53,58,59). Some reports indicate that CBP also serves as an acetyl transferase for several DNA repair proteins, such as Ku70, PCNA, TDG, while both p300 and CBP acetylate tumor suppressor, p53 (59). In this study, we show that PNKP, involved in multiple DNA repair pathways, is acetylated by both acetyl transferases, p300 and CBP, at two distinct lysine residues. It was found that p300 acetylates K142 constitutively and CBP acetylates at K226 only after DSB induction. We have previously shown that NEIL1 and NEIL2 stably associate with p300, forming specific BER complexes along with PNKP (49,50,53). The association of AcK142 with TC-BER/SSBR proteins, along with the elongating form of RNA polymerase II and specifically with p300, is therefore consistent with our previous reports. Interestingly, we found the presence of a NHEJ protein Ku70 in the Ac-K142 PNKP immunocomplex. This finding is corroborative with a recent report that shows BER-NHEJ cross-talk is mediated by the formation of a Ku70-Polβ complex at the repair foci and impairment of BER efficiency occurs via ablation of Ku70 (68). On the other hand, we found an association of K226 with CBP, RNAP II and NHEJ factors following DSB induction. To our knowledge, this is the first example of acetylation of a DNA repair protein at two distinct sites by two different acetyl transferases, where such post-translational modifications of PNKP act as a critical determinant of its role in SSBR vs. NHEJ. Notably, AcK226 PNKP immunocomplex showed the presence of DNA polymerase eta (Pol η). We recently provided evidence that Pol η plays a critical role in copying the sequence information via its reverse transcriptase activity from nascent RNA into DNA at the DSB sites (19). Thus, we postulate a role of Ac-K226 PNKP in the recruitment of Pol η for RNA-templated error-free repair of DSBs via the TC-NHEJ pathway.

Mutations in PNKP and resulting repair deficiency have been implicated in a variety of human neurological diseases, such as MCSZ, AOA4, etc (28,60–63). In our previous study, we also demonstrated that PNKP activity, not its protein level, is severely abrogated in Huntington’s disease (HD), a major polyglutamine disease, resulting accumulation of DSBs (18,24). In the present study, we found that acetylation at K226, but not at K142, is impaired in mouse Huntington’s Disease (HD) derived striatal neuronal cells (Q111), which is consistent with previous reports and our present study showing selective degradation of CBP in HD mouse-derived striatal neuronal cells (Q111) compared to WT (Q7) cells. Consequently, Q111 cells showed progressive accumulation of DSBs in the transcribed genes, which can trigger apoptosis to vulnerable brain cells: a plausible cause of the neurodegeneration in HD. In summary, we have provided critical evidence for the role of acetylation at two distinct residues located in different domains of PNKP that regulate its functionally distinct roles in TC-BER/SSBR vs. TC-NHEJ in mammalian cells.

## Acknowledgments

This work was supported by National Institute of Health Grants 2R01 NS073976 to TH; R01HL145477 to TH and Sanjiv Sur (SS), Division of Allergy and Clinical Immunology, Baylor College of Medicine, Houston; National Institute of Allergic and Infectious Diseases (NIAID) grant AI062885 to IB; and University of California Tobacco-Related Disease Research Program (TRDRP) grant 26IR-0017 to A.H.S. We thank Dr. Katherine Kaus, Research Development Specialist at the University of Texas Medical Branch for editing this manuscript.

## Author contributions

TH conceived the research. TH, AI and AC designed the research. AI, AC, AS and UA performed research. IB and GS provided reagents and critical intellectual input. AI, AC and TH wrote the manuscript. All the authors read and approved the final version of the manuscript.

## Data availability

The data that support the findings of this study are available from the corresponding author upon request.

## Supplementary Figures

**Supplementary** Figure 1. Identification of two novel acetyl lysine sites in PNKP by LC-MS/MS. (A) Acetylation sites are marked in red within the peptide sequence and the amino acid residues are listed in a subtitle as well. The table shows the Sequest algorithm scores (Xcorr and ΔCorr) along with the mass accuracy measurement of the acetylated peptides. (B and C) The figures show the fragmentation pattern of two acetylated peptides matched to PNKP. Representative figures of acetyl-lysine spectrums of PNKP are shown: (B) the spectrum of peptide sequence for K142 acetylation and (C) the spectrum of peptide sequence for K226 acetylation. The acetylated lysines are shown as K#. The # symbol represents a mass addition of 42.0106 Da to lysine. (D) Schematic representation of PNKP domains (FHA, Linker, phosphatase and kinase), indicating the K142 acetylation site in the linker region and the K226 acetylation site in the phosphatase domain. (E) Results of a similar Mass Spectrometric analysis are represented in the table showing the acetylated residues in WT vs. K226R PNKP stable cell lines ±Bleo.

**Supplementary** Figure 2. (A) Western blot shows the expression level of phosphorylated 53BP1, total 53BP1 and γH2AX in the chromatin fraction of Bleo-treated WT (lane 1) and K226R (lane 2) and GO-treated WT (lane 3) and K142R (lane 4) cells. HDAC2: used as a loading control. (B) Depletion of endogenous PNKP by 3’-UTR specific siRNA. The representative agarose gel shows the extent of depletion of endogenous PNKP in WT, K142R, K226R and K142R/226R PNKP expressing stable cell lines by 3’UTR specific siRNA. GAPDH was used as housekeeping control (upper panels). The bar diagram represents the relative expression level of endogenous PNKP normalized with the expression of control GAPDH with the control siRNA transfected samples considered as 100 (Error bars represent ±SD of the mean, n=3, ***P<0.005) (lower panel).

**Supplementary** Figure 3. Depletion of endogenous p300 and CBP by siRNA in HEK293 cells. Western blots show the relative expression level of (A) p300 and (B) CBP in the chromatin fraction following control (Con; lane 1) and specific siRNA (lane 2) transfection. HDAC2 is used as a loading control.

**Supplementary** Figure 4. Depletion of endogenous PNKP by 3’-UTR specific siRNA. The representative agarose gels (upper panels) show the extent of depletion of endogenous PNKP in WT, K142R, K226R and K142R/226R PNKP expressing stable cell lines (A) and WT, K142Q, K226Q and K142Q/226Q PNKP expressing stable cell lines (B) by 3’UTR specific siRNA. The bar diagrams represent the relative expression level of endogenous PNKP normalized with the expression of control GAPDH with the control siRNA transfected samples considered as 100 (Error bars represent ±SD of the mean, n=3, ***P<0.005) (lower panels).

**Supplementary** Figure 5. (A and C) (Upper panels) The representative agarose gels show the extent of depletion of endogenous PNKP in (A) WT, K142R, K226R and K142R/226R PNKP expressing stable cell lines and in (C) WT, K142Q, K226Q and K142Q/226Q PNKP expressing stable cell lines under the same experimental conditions. (Lower panels) The bar diagrams represent the relative expression level of endogenous PNKP normalized with the expression of control GAPDH with the control siRNA transfected samples considered as 100 (Error bars represent ±SD of the mean, n=3, ***P<0.005). (B and D) Western blots show the expression of γH2AX (upper panels) in the chromatin fraction of (B) WT, K142R, K226R and K142R/226R PNKP expressing cells and (D) WT, K142Q, K226Q and K142Q/226Q expressing cells mock (C), Bleo-treated (B) or 16 h post Bleo-treatment (R). HDAC2 is used as the loading control (lower panels).

