## Supplementary Figures for "Site-specific acetylation of polynucleotide kinase 3’-phosphatase (PNKP) regulates its distinct role in DNA repair pathways"

Supplementary Figure 1

(A)

| PPM | XCorr | ΔCorr | Peptide | Target amino acid |
|---|---|---|---|---|
| 0.79 | 3.422 | 0.555 | R.**K#**SNPGWENLEK.L | K142 |
| 0.92 | 2.091 | 0.694 | R.G**K#**LPAEEFK.A | K226 |

(B)

Predicted Fragmentation Pattern

| Seq | # | b: Δ Error | b | y | y: Δ Error | +1 |
|---|---|---|---|---|---|---|
| K# | 1 | --- | 171.1 | --- | --- | 11 |
| S | 2 | -0.1 | 258.1 | 1173.6 | 0.1 | 10 |
| N | 3 | 0.0 | 372.2 | 1086.5 | 0.0 | 9 |
| P | 4 | 0.2 | 469.2 | 972.5 | 0.0 | 8 |
| G | 5 | 0.0 | 526.3 | 875.4 | 0.0 | 7 |
| W | 6 | 0.0 | 712.3 | 818.4 | 0.1 | 6 |
| E | 7 | 0.0 | 841.4 | 632.3 | 0.0 | 5 |
| N | 8 | 0.3 | 955.4 | 503.3 | 0.0 | 4 |
| L | 9 | -0.1 | 1068.5 | 389.2 | 0.1 | 3 |
| E | 10 | 0.0 | 1197.6 | 276.2 | 0.2 | 2 |
| K | 11 | --- | --- | 147.1 | --- | 1 |

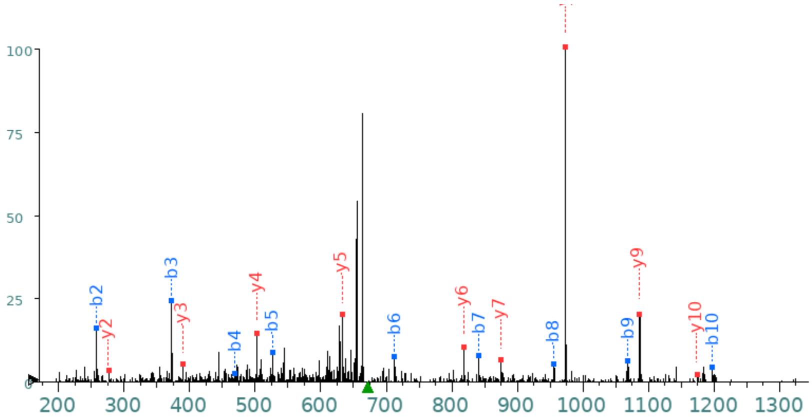


(C)

Predicted Fragmentation Pattern

| Seq | # | b: Δ Error | b | y | y: Δ Error | +1 |
|---|---|---|---|---|---|---|
| G | 1 | --- | 58.0 | --- | --- | 9 |
| K# | 2 | 0.1 | 228.1 | 1003.5 | --- | 8 |
| L | 3 | 0.0 | 341.2 | 833.4 | 0.2 | 7 |
| P | 4 | 0.0 | 438.3 | 720.4 | 0.0 | 6 |
| A | 5 | 0.0 | 509.3 | 623.3 | 0.1 | 5 |
| E | 6 | 0.2 | 638.4 | 552.3 | 0.3 | 4 |
| E | 7 | 0.0 | 767.4 | 423.2 | 0.1 | 3 |
| F | 8 | 0.1 | 914.5 | 294.2 | 0.1 | 2 |
| K | 9 | --- | --- | 147.1 | --- | 1 |

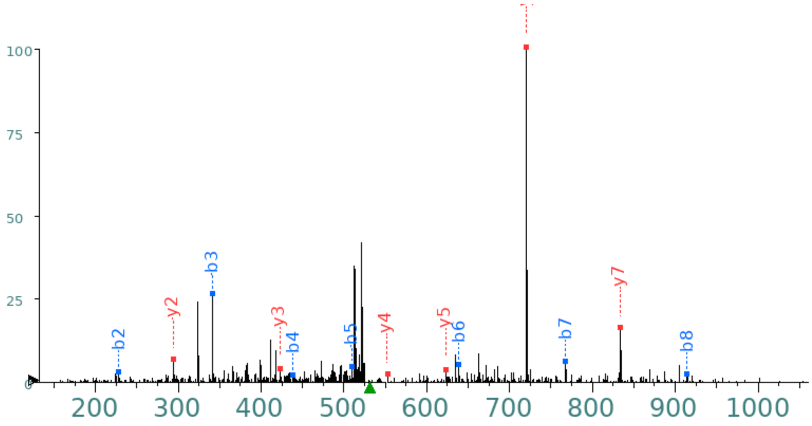

Supplementary Figure 1 (contd)

(D)

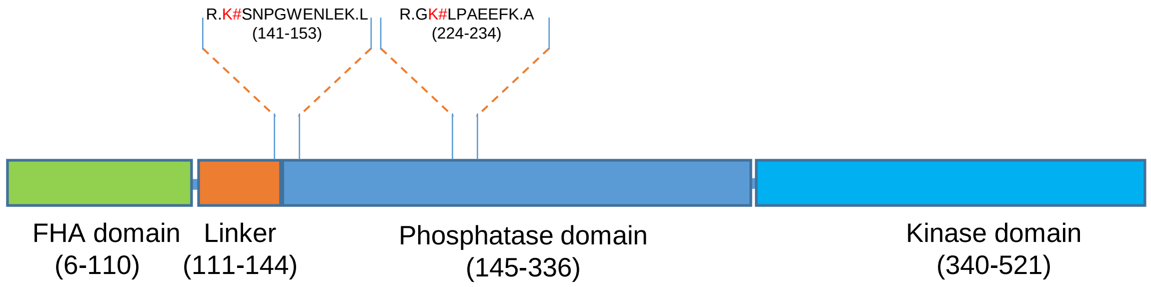


(E)

| PNKP | Treatment | Acetylated amino acid # |
|---|---|---|
| WT-FLAG | Mock | K142 |
| WT-FLAG | (+) Bleo | K142/K226 |
| K226R-FLAG | Mock | K142 |
| K226R-FLAG | (+) Bleo | K142 |

Supplementary Figure 2

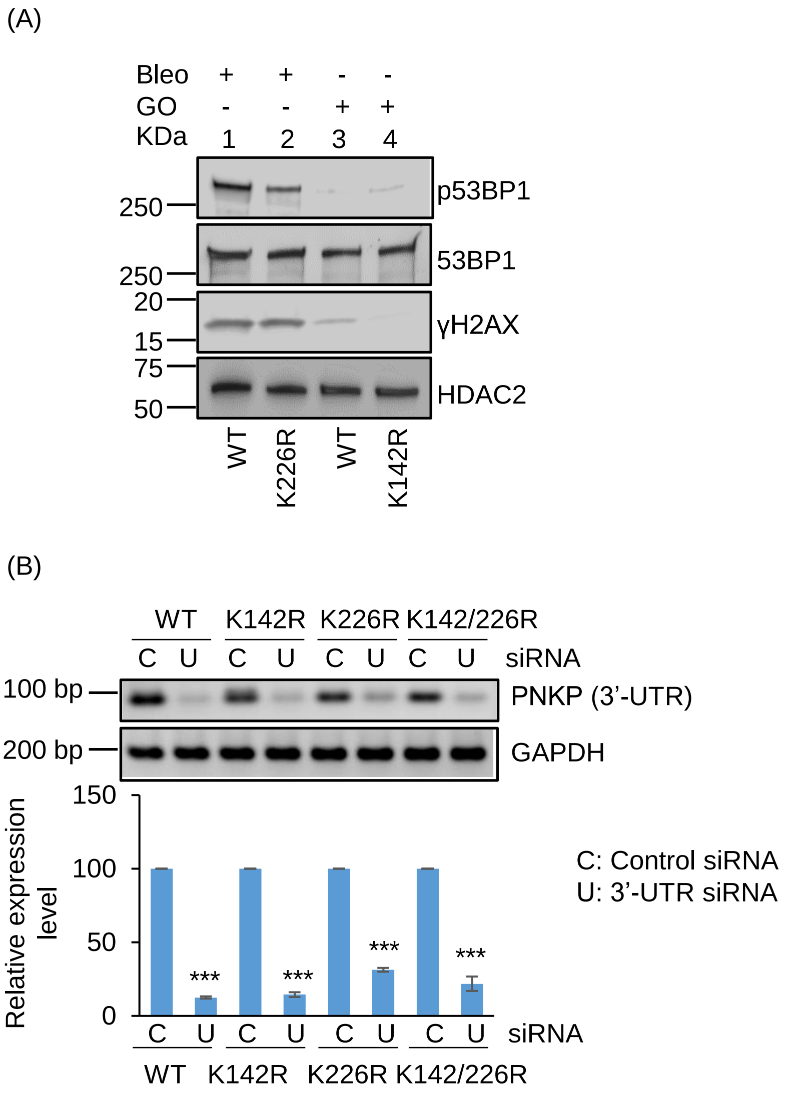

Supplementary Figure 3

(A)

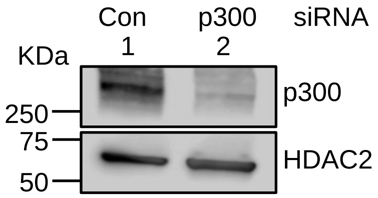


(B)

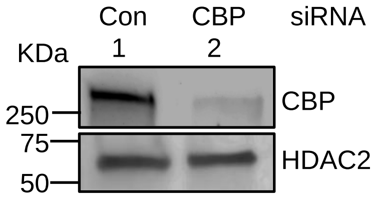

Supplementary Figure 4

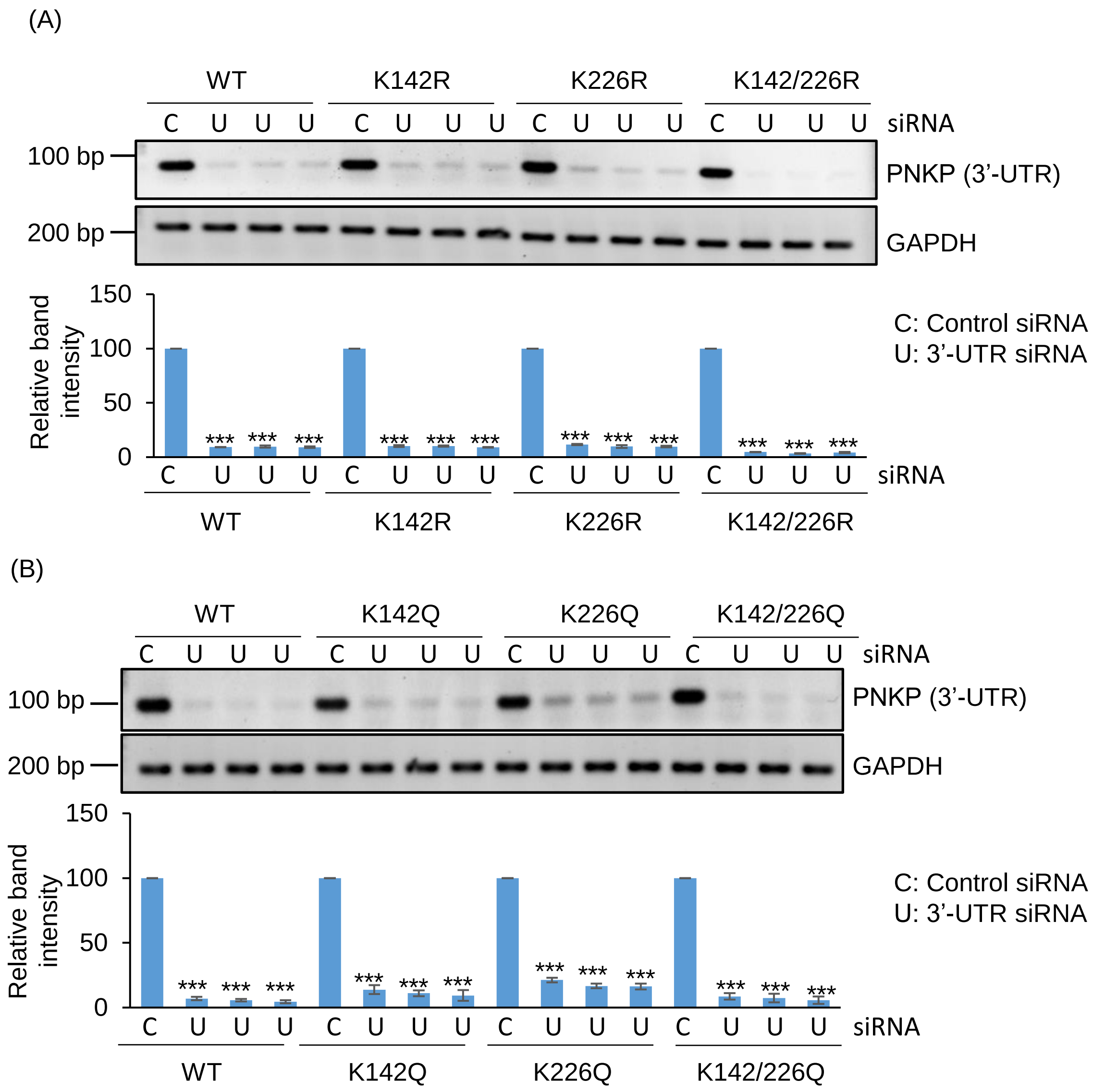

Supplementary Figure 5

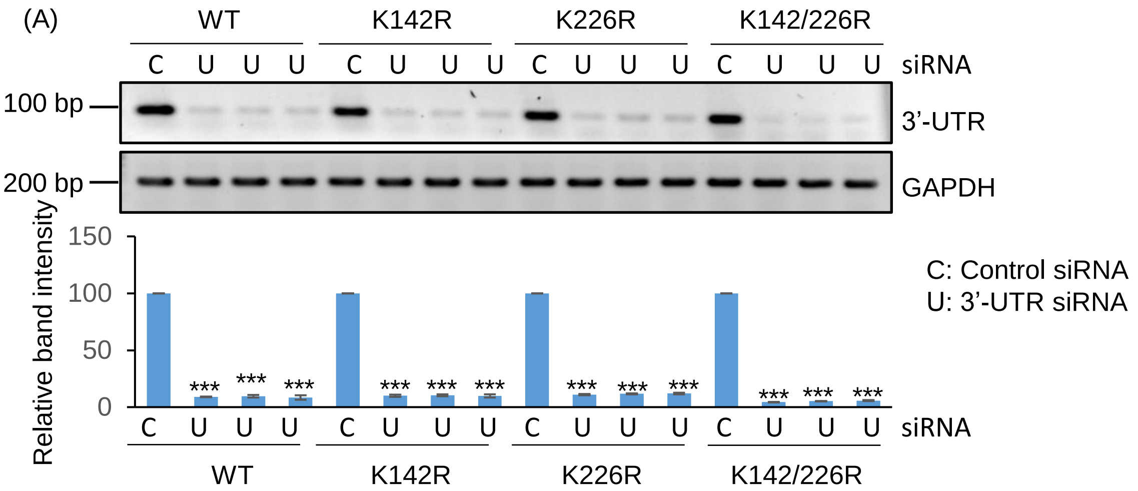


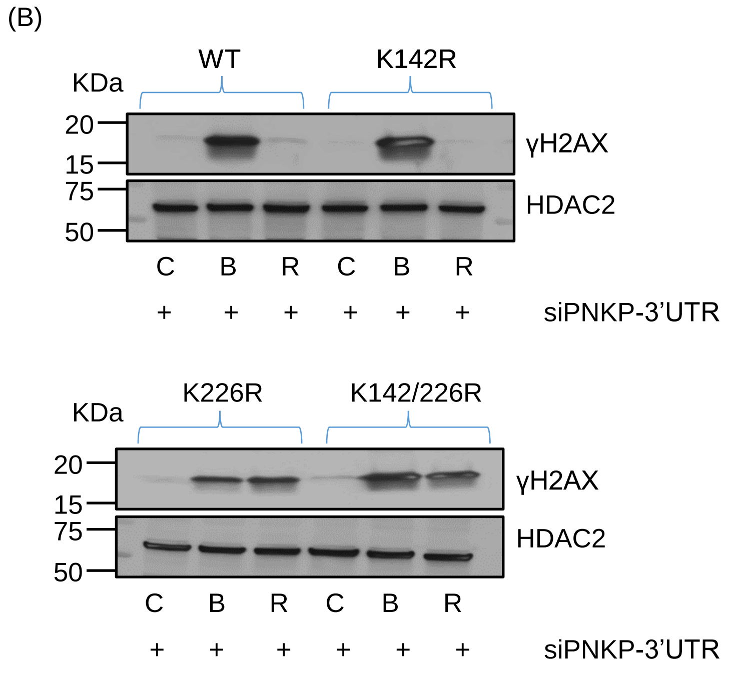

Supplementary Figure 5 (contd)

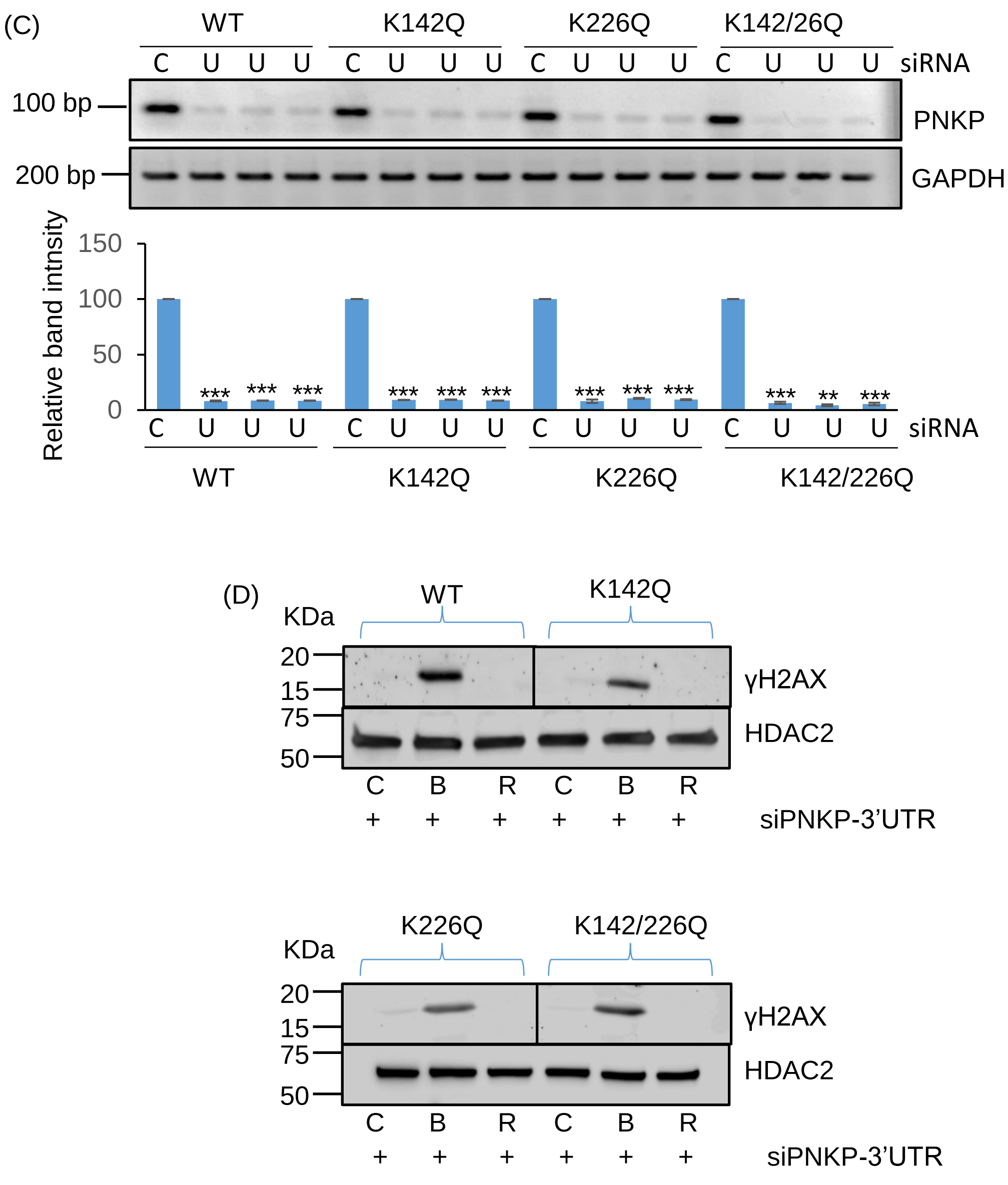
